# On-chip light diffraction imaging of nano structures in the guanine platelet

**DOI:** 10.1101/2021.08.15.456435

**Authors:** Masakazu Iwasaka

**Affiliations:** Hiroshima University, Kagamiyama 1-4-2 Higashihiroshima, Hiroshima 739-8527, Japan

## Abstract

Light projection over short distances can minimize the size of photonic devices, e.g., head-mounted displays and lens-free microscopes. Small lenses or light condensers without typical lenses are essential for light control in micron-scale spaces. In this work, micro-platelets floating in water are used for light projection near the image sensor. These platelets, which are made from guanine, have nanohole gratings and demonstrate light diffraction toward specific directions. By setting a thin water layer on the image sensor’s cover glass, each platelet in water forms column-or bar-code-shaped images on the screen. The projected image shapes and colors are inferred to contain information about nano-structures present in the guanine platelet. The proposed down-sized imaging technique can realize extremely compact and portable imagers for nanoscale object detection.

---

The miniaturization of photonic devices is contributing to the diversity of the ubiquitous technology available in modern society. In portable devices such as mobile phone and small life-science-oriented lens-free microscopes, charge-coupled devices (CCD) or complementary metal-oxide-semiconductor (CMOS) image sensors are widely used. The latter sensors are used to perform on-chip imaging (1-3), which offers the advantage of realizing imaging of cells and living tissues within a compact space (4-12).

The development of endoscopes and cell culture microscopes without lenses reduced the size of the apparatus required and its performance was improved by applying computational analysis. In the case of application to cellular imaging, microfluidic devices were shown to be effective in locating targeted cells close to the image sensor. For the illumination optics, the so-called shadow imaging method was applied with further image processing. Image reconstruction using holographic images and a deep learning approach (11,12) were used to address the challenge of obtaining the same spatial resolution that is provided by a microscope with a lens.

However, additional potential remains for further miniaturization of the illumination optics. Utilization of planar films with nano-structures such as metasurfaces offers one potential approach for reducing the size of the illumination optics. (13, 14) Various types of flat optics structures were reported to act as functional lenses, thus achieving focusing without using a classical lens. (15-16) Nano-structures such as nanoholes (17-21) or grating (22) formed in a substrate have also achieved light focusing abilities.

Over the last two decades, studies of artificial nano-fabrication techniques for flat optics have opened up new ways to obtain flat lenses. In contrast, with regard to lenses in living systems, studies have continued to focus on classical dielectric lenses. There have been no examples of biological flat optics using similar mechanisms to those reported recently in optical material science.

In the bio-photonics field, as a material responsible for control of biological light reflection, guanine platelets obtained from teleost fish skin have been investigated since the 1960s (23). The light reflecting properties of these platelets were explained (24) by considering the platelet to be a thin uniform platelet composed of guanine, which is one of the nucleic acid base molecules. Recently, scanning electron microscopy (SEM) and transmission electron microscopy (TEM) analyses of the guanine platelets of two species revealed that these platelets have sub-micrometer structures composed of triangular holes in each platelet (25) that were formed during or after their bio-crystallization process.

Figure 1 shows that optical microscopes can also visualize these sub-micrometer structures by tracking slight changes in the illumination conditions. In Fig. 1A, observation of the grating on the main face of a guanine platelet was achieved during a slight tilting motion of the platelet. Figure 1 and Figures S1-S4 indicate the existence of a grating with a width of less than 1 μm. During slight tilting of the tiny, thin platelet, images indicating the existence of nano-holes in the platelet were acquired. The platelet shown in Fig. 1B had holes arranged along the sides of triangular shapes, which corresponded to the triangular structure observed in the corresponding SEM image (Fig. 1D).

**Figure 1.**
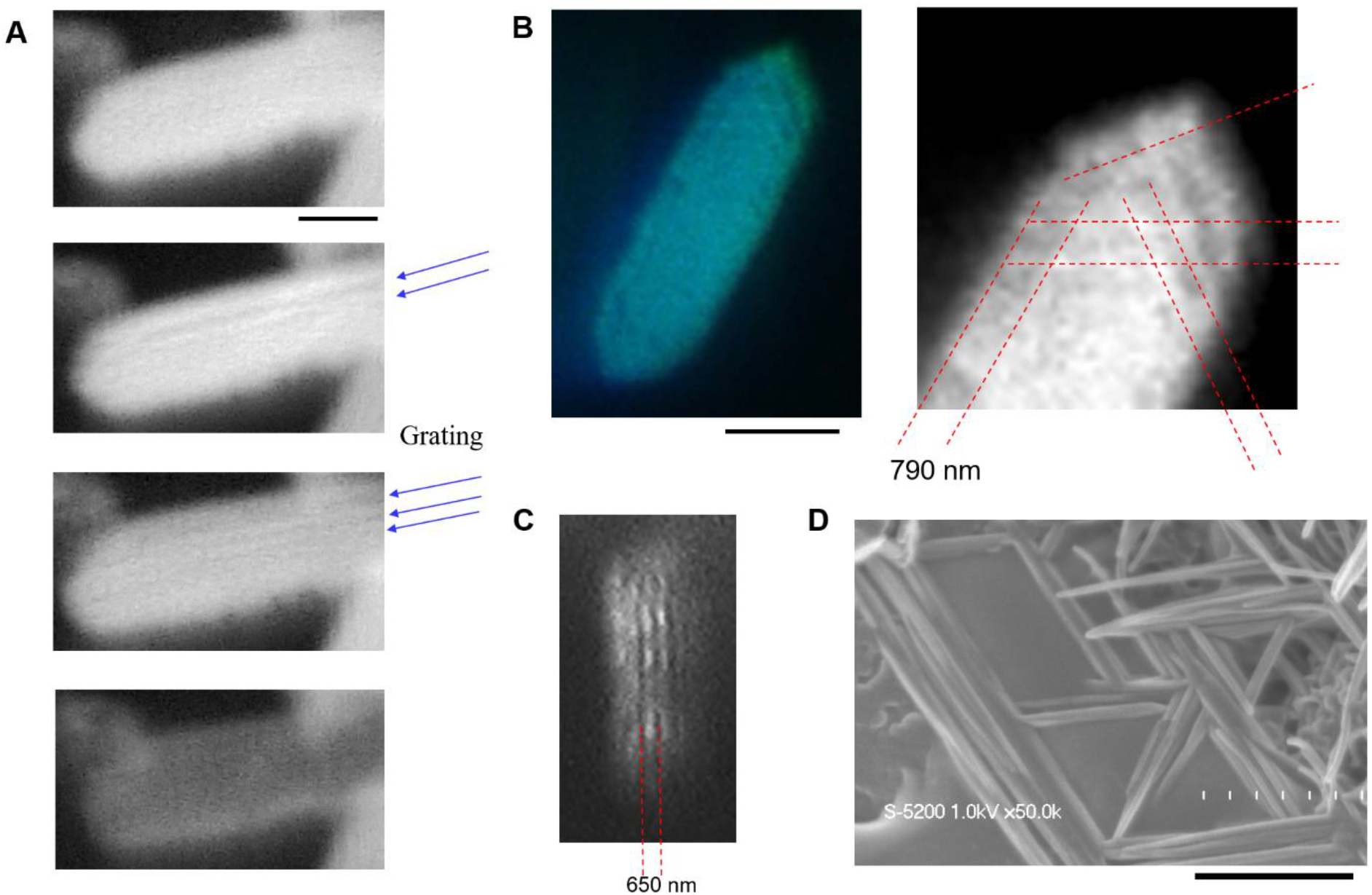
Grating structure observed in microscope images of fish guanine platelets. (A) Optical microscope images of a guanine platelet (main face) from a goldfish, *Carassius auratus*. The scale bar represents 5 μm. (B),-(C) Optical microscope images of guanine platelets from a Japanese anchovy, *Engraulis japonicus*. The scale bar represents 4 μm. (D) SEM image of the Japanese anchovy platelet. The scale bar represents 600 nm.

It was found that these fish guanine platelets in water caused strong contrast images to be observed on a CMOS image sensor array without a lens (26, submitted). In the improved imaging system shown in Fig. 2 (and Fig. S5), on-chip imaging was performed by directing the sensor array downward toward a black substrate, which acted to provide light shading. The length of gap between the image sensor and the black substrate was adjustable from 0.5 mm up to 2 mm. A water layer containing the guanine platelets was then formed in this gap. The tilting angle of the guanine platelets in water was uncontrolled, which meant that various types of light scattering occurred on both the reflection side and the transmission side (Fig. 2A).

**Figure 2.**
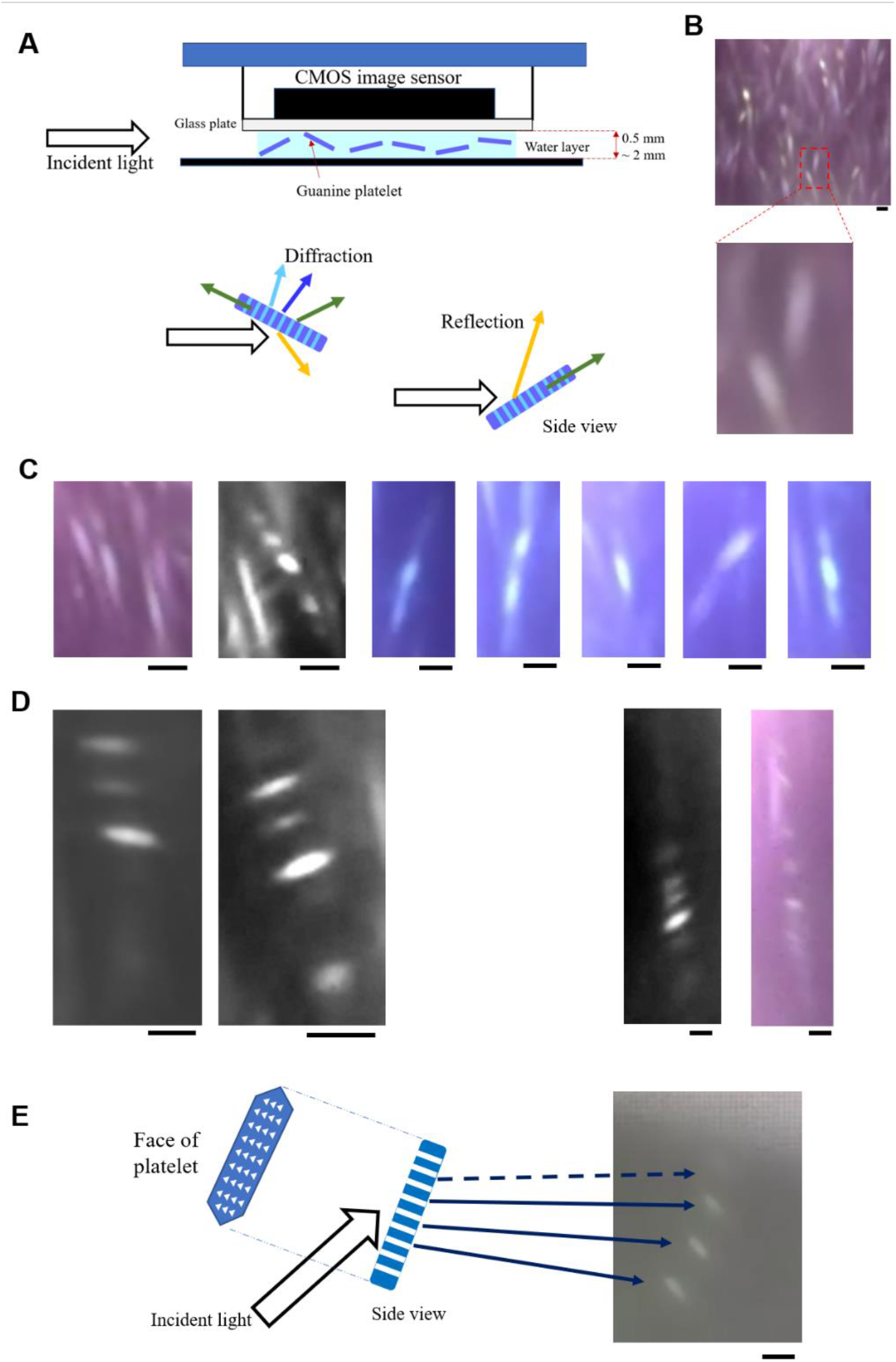
On-chip light scattering imaging of tiny goldfish platelets. (A) Scheme for imaging of platelets floating in water that form a thin layer beneath a CMOS sensor chip. The lower part of the illustration shows the speculated light scattering patterns. (B)- (D) Typical image patterns projected on the sensor array without the lens, where (B) shows the platelet-like pattern, (C) shows the column-type pattern, and (D) shows the bar-code-type pattern. (E) Speculated relationship between a platelet and projected images. The scale bar represents 100 μm.

When the gap thickness was varied, column-shaped or bar-code-like images were obtained (Fig. 2C and 2D, respectively), along with the platelet-like projection image (Fig. 2B) on the image sensor (*see* Supplementary movie S1). It was thus possible, that the tiny platelets with sub-micrometer-scale holes contributed to the arrangements of the light scattering patterns appearing on the reflection side and the transmission side (Fig. 2E).

Apart from the three typical image types (Fig. 2B-D and Fig. S6-S10), other variant images appeared as an incidental case (see Fig. 3 and Fig. S11-12). These colored, large-scale or assembled patterns involved column-type or bar-code-type components (*see* Supplementary movie S2). When compared with the typical patterns, these patterns were larger in size and there was a clear occurrence of light dispersion (Fig. 3A). The two large patterns that appear in Fig. 3A (top panel) individually caused a to-and-fro motion. This indicates that the projection originated from one object. These patterns were provably composed of complexes of platelets, such as stacked platelets. Because the platelets were obtained from fish scales, they are likely to be contaminated with biological substances (e.g., protein, lipids). Although, the variants shown in Fig. 2B were not colored, the patterns were composed of eight to nine bar-code shapes that moved together. A complex composed of two to three platelets floating in the water provably projected the patterns onto the image sensor plane.

**Figure 3.**
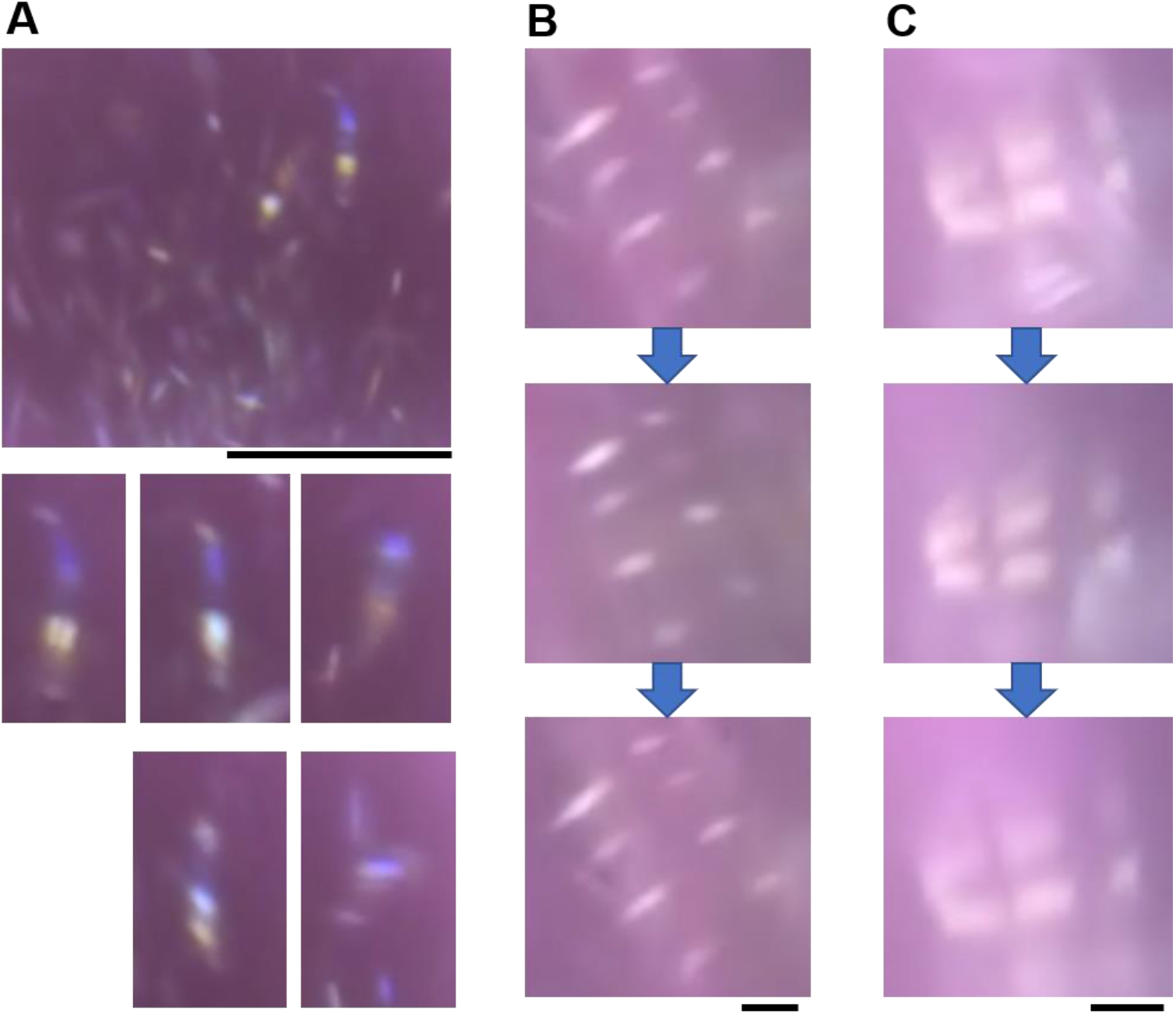
Variant light scattering patterns produced by guanine platelets from Japanese anchovy. A) Colored patterns appearing in a mass of images (top). The scale bar represents 1 mm. The middle and bottom panels show five examples of the colored light scattering images. B) Light scattering patterns from two connected platelets (left) and three connected platelets (right). The arrows indicate the time series of the changes in the image. The scale bar represents 100 μm.

To clarify the light scattering patterns from the guanine platelets observed in the lens-free CMOS sensor array, a numerical simulation was performed using the finite-difference time-domain (FDTD) method (Fig. 4 and Fig. S13-22). Specifically, it was impossible to explain the generation of the bar-code pattern using regular reflections only. In fact, regular reflections (and forward scattering in the direction parallel to that of the incident light) provide the maximum intensity for the scattered light. It is thus necessary to find a comparable sub-branch among the scattering patterns. The dependence of the formation of the scattering pattern (logarithmic spherical plot) on the angle of incidence, the wavelength of the incident light, and the presence of the grating was thus investigated (Fig. 4A-C).

**Figure 4.**
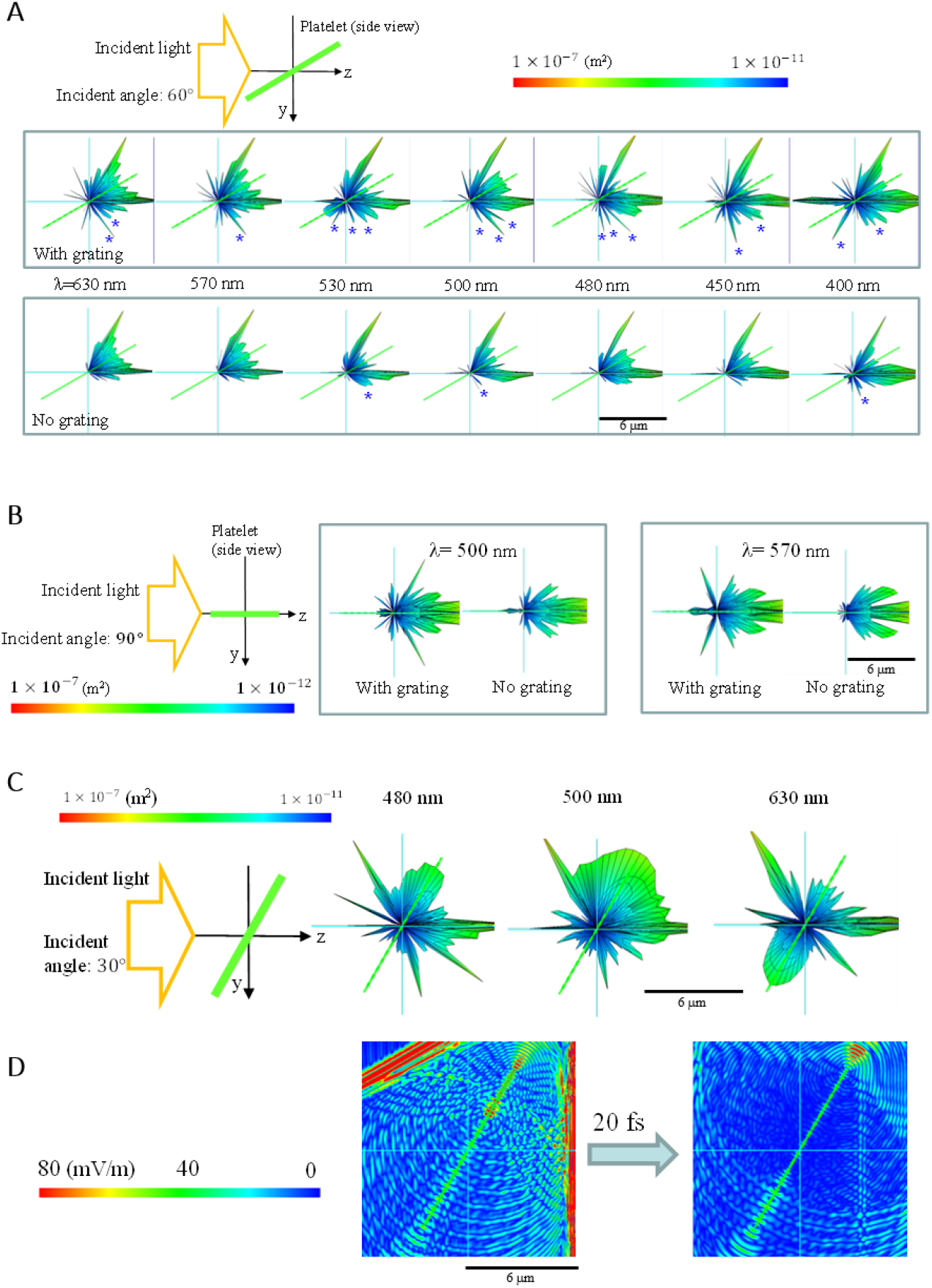
Electromagnetic FDTD simulation results of light scattering by a guanine platelet floating in water. The platelet contained a grating formed by an alignment of nano-holes filled with water. The refractive index of guanine was set at 1.83. (*see* the Supplementary Material for more detailed specifications of the simulation model). The incident light propagating along the z-axis was *s*-polarized when the incident plane was set to be the y-z plane. A- C show spherical plots of the scattered light over the wavelength range from 630 nm to 400 nm. (A) Comparison of light scattering by a guanine platelet with and without a grating when the angle of incidence was 60°. The blue asterisk marks a distinct branch of the scattered light appearing on the transmission side. (B) Light scattering by a guanine platelet when the angle of incidence was 90° (for λ = 500 nm and 570 nm). (C) Light scattering by a guanine platelet when the angle of incidence was 30° (only for the platelet with the grating at λ = 480 nm, 500 nm and 630 nm). (D) Electric field distribution around the platelet after the incident light passed through it. The color indicators indicate the cross-sectional area (m^2^) in (A)- (C) and the electric field in (D). The scale bar represents 6 μm.

In Fig. 4A, the angle of incidence with respect to the main face (i.e., the broadest plane) of the platelet was 60°. When compared with a platelet without the grating (bottom panel of Fig. 4A), the platelet with the grating (middle panel of Fig. 4A) had more light scattering branches in the transmission side. Figure 4B shows the case where the main face of the platelet was oriented to be parallel to the incident light. It is obvious that the grating in the platelet was inducing the inclination in the vector of the light scattering normal to the main face. Strongly and highly-directionally scattered light was directed at angles of less than ±30° with respect to the direction normal to that of the incident light. This tendency became more distinct when the angle of incidence was 30° (Fig. 4C). The scattering pattern was symmetrical, with reflection peaks and diffraction peaks existing in equal numbers. In addition to the reflection and diffraction peaks, a spherical plot of the scattered light in Fig. 4c shows that strong light (radiation-like) scattering occurred in the direction parallel to the platelet surface for light at wavelength of 500 nm and 630 nm. The intensity of this strong and broad scattering behavior was similar to that of regular reflections. Figure 4D indicates that the unique scattering that occurred parallel to the platelet was related to the light propagation inside the 110-nm-thick platelet. The internal light propagation continued for 20 fs after the wave-front of the incident light passed through the platelet. The distances between the holes in the platelet simulation model were 540 nm and 270 nm, as shown in the Supplementary Material. The light scattering pattern can be varied by changing the geometrical parameters of the platelet and the grating (holes). A platelet without a grating can also generate the diffraction branch in the light transmission side of the platelet. However, formation of the hole grating apparently enhanced the intensity of the diffractions, which were related to the bar-code-type projection on the CMOS sensor.

The results indicate that the fish guanine platelets used in this case acted as a diffractive lens (27). Unlike an artificial diffractive lens formed in a film on a substrate, the biogenic fish guanine platelets used in these experiments were initially formed in the organelles of a living cell and were then extracted and dispersed in an aqueous solution. Because the platelets can be floated individually in water, they can operate as a three-dimensional diffractive lens in a thin water layer.

The three basic three types of scattering pattern can be attributed to the structure of the individual platelets and the tilting angle versus the angle of light incidence. These simple image signals from the tiny platelets floating in water are suitable for detection of micro-environmental conditions. Imaging of both micromechanical motion and micro/nanoscale structures is possible when the guanine platelets are used as an on-chip reflection imaging transducer. We can make the angle of incidence higher, reduce the thickness of the illumination optics, and then use the advantages of the tiny, thin platelets, which not only offer strong reflection but also have diffraction lens properties. Other holographic-type on-chip imaging methods have also used the light diffraction patterns of the targeted object when the incident light was provided along the optical axis. In the on-chip imaging process in this work, only the fish guanine platelets exhibited images with strong contrast, while other particles suspended in the water did not show images with the same contrast properties.

If the colorful variants shown in Fig. 3 were the product of stacked platelets, the variant image should then be a consequence of the interactions between the tiny platelets and other objects. A component with a bar-code pattern originated from a guanine platelet with dimensions of 20- 40 μm × 5- 10 μm × 100- 160 nm. Therefore, the emergence of the variants (as shown in Fig. 3B) indicates integration of the platelets. For future applications of on-chip light diffraction imaging with fish guanine platelets, use of additional image analysis methods such as deep learning techniques may be effective.

## Acknowledgments

The author thanks David MacDonald, MSc, from Edanz (https://jp.edanz.com/ac) for editing a draft of this manuscript.

## Author contributions

MI performed the experimental design, experiments, measurements and analyses. All parts of the manuscript and illustrations were prepared by MI.

## Data and materials availability

All data is available in the main text or the supplementary materials.

