## Supplementary Materials for "On-chip light diffraction imaging of nano structures in the guanine platelet"

M. Iwasaka[1]*

[1]Hiroshima University; Kagamiyama 1-4-2 Higashihiroshima, Hiroshima 739-8527, Japan.

**Materials and Methods**

Guanine platelets of fish

The experimental methods used with the fish were approved by Hiroshima University Animal Care and Use Committee (approval numbers F16–2 and F20-4, Hiroshima University). The guanine platelets were obtained from the scales and skins of the goldfish, *Carassius auratus* and the Japanese anchovy, *Engraulis japonicus*. An aqueous suspension of the guanine platelets ($10^4$-$10^5$ per ml) was prepared using distilled water. The purification protocol used for the guanine platelets in water was the same as the methods that was reported previously in (28).

Optical microscope (with lens) and scanning electron microscope (SEM)

The optical images shown in Fig. 1 were acquired using a high-resolution microscope lens (MXB-2500REZ; Hirox, Tokyo Japan). The optical microscopic observation system (RH-2000; Hirox, Tokyo, Japan) was used. The SEM images were acquired by using a field-emission scanning electron microscope (FE-SEM; S-5200, Hitachi, Tokyo, Japan) (25). Figure S1 shows two types of optical observation method. Provision of light through the inner light guide of the lens (Fig. S1B) brought a bright field image shown in Fig. S2. The image of the guanine platelet, which was separated from a goldfish scale, showed cleavages on its surface. The observed cleavage lines were aligned in parallel with the edge of the platelet. This cleavage pattern geometry is consistent with the images obtained in a previous study performed on the SEM (26).

The observation method shown in Fig. 1A uses asymmetric illumination from a specific direction provided by a separate illumination unit (optical fiber-guided light). By using this method, which is a type of dark field illumination, we can increase the probability of observation of the nano-grating structure on a guanine platelet, as illustrated in Figs. S3 to S5.

The guanine platelets were observed in water that was sandwiched between a pair of glass plates. These platelets were floating in the water or were partially attached to the surface of one of the glass plates. The photographs shown in Figs. S3 to S5 indicate that the observed platelets changed their surface textures when they were tilted slightly. The platelets appeared to have smooth surfaces when they reflected light strongly. These examples of the platelet showed a surface with gratings that depended on the platelet's tilt angle.

CMOS image sensor chip for lens-free imaging

As the image sensor used in the experiments, a CMOS module (USB camera module ELP-SUSB1080PO1-L36, Ailipu Technology, Shenzhen, Guangdong, China) was used after its lens was detached. As shown in Fig. S6, the image sensor chip was mounted on the beam of a stand with a revolution capability that enabled its height adjustment. 50 μl of an aqueous suspension containing the guanine platelets was placed on a black metal plate (sample stage; top right panel of Fig. S6). Closing the glass plate that covered the image sensor onto the aqueous suspension on the plate from top to bottom formed a thin water layer between the image sensor and the sample stage.

Illumination and sensor chip approach to the sample for the lens-free imaging

A sample stage was placed behind the image sensor to direct its glass-covered surface downward. After 50 μl of the guanine platelet suspension was deposited on the surface of the stage (black colored metal plate), the glass plate covering the sensor was then closed to the aqueous suspension and a layer of water was formed between the two plates. By adjusting the image sensor chip height, the gap for the water layer was varied manually from approximately 0.5 mm to 2 mm. The incident light, which was provided by a light-emitting diode LED source and a light guide, was introduced into this gap as shown in Fig. S6. The angle between the incident light and the optical axis in the conventional case was nearly 90°. The LED source used here was the LA-HDF158A (Hayashi Repic, Tokyo, Japan). The diameter of the light guide was approximately 8 mm.

Figures S7 and S8 present examples used to testing how the image of the guanine platelets that was projected directly varied when the width of the water layer containing the floating guanine platelets changed.

Image analysis for lens-free imaging

The movies of the lens-free imaging behavior of the floating guanine platelets were collected using a personal computer to which the CMOS imaging module was connected. The movie recordings were performed using image capture software (Xploview, VIXEN, Saitama, Japan), in which the white balance was set manually at 4650 K. Static images were then captured from playback of the recorded movies. Images in the region of interest (ROI) were selected using Image J software (1.50i, NIH, USA).

Figures S9- S12 show examples of the image capture and selection processes, although these examples were not used to form the presentations shown in Fig. 2 and Fig. 3 in the main text. After adjustment of both the water layer thickness and the light illumination direction, the movie recording started. The maximum width of the entire captured image area was approximately 4 mm. As noted in the main text, the directly projected floating platelets patterns on the image sensor were categorized into individual types with general or specific patterns. The ROI was determined based on the purpose of the selection, i.e., obtaining images of general (platelet-, column-, or bar-code shaped) patterns or variant patterns.

Figures S9 and S10 show evidence that the bar-code images moved together. The bar-code patterns were observed frequently when the water layer thickness increased once after having decreased. The images shown in Figs. S9 and S10 were selected by searching the bar-code patterns for shifting (or drifting) without changes in the distances between the bars during the movie playback. Typically, three to four bars were observed to cause a to-and-fro motion together. In the case shown in the upper panel of Fig. S10, seven bars were found to cause this type of motion. The image behaved like a long ladder.

Figures S11 and S12 present complementary data as evidence that a few colored images or complex images appeared among the mass of projected images.

Numerical simulation (FDTD)

The light scattering patterns around a guanine platelet floating in water were simulated numerically. A commercial electromagnetic field solution software package, Poynting for Optics V03L10R121 (Fujitsu, Tokyo, Japan) was used. This software uses the finite-difference time-domain (FDTD) simulation algorithm, which enables visualization of both the light scattering pattern and the electric field distribution around the guanine platelet floating in water.

In the model (as shown in Fig. S13), a pulsed light beam propagated toward the +*z* direction. The incident light had a rectangular volume, i.e. it was from a three-dimensional light source. Each orange-colored arrow in Figs. S13 to S22 indicates the direction of the incident light that propagated inside a rectangular box filled with water (refractive index $n$ = 1.33). After the near-field electromagnetic fields of the light propagating around the platelet were calculated, a spherical plot of the scattered light was then calculated. Because the pulsed light involved frequency elements from the visible light range (as shown in the light source shape in Fig. S13), we can obtain the light scattering intensities at 400 nm, 450 nm, 480 nm, 500 nm, 530 nm, 570 nm, and 630 nm.

In the case of a mono-plate (i.e., with no holes), the geometry of the guanine platelet, for which the refractive index $n$ = 1.83, was set at 5 μm × 8 μm for the main face dimensions and 110 nm for the thickness. Model platelets with nano holes are shown in Figs. S13 and S19. Figure S13 shows the model used for the presentation in the main text, which consisted of triangular holes penetrating the platelet. The refractive index inside each hole was 1.33 (i.e., that of water). The holes aligned with cycles of 270 nm (along the *x*-axis) and 540 nm (along the *y*-axis). This alignment was determined based on consideration of the SEM images obtained (as shown in Fig. 1 and in reference (26)). The angle of incidence was varied by tilting the platelet without changing the light source direction. In Fig. S13C, where the platelet was oriented to be parallel with the incident light, the angle of incidence was 90°. Figure S13D shows the model obtained when the angle of incidence was 30°.

The spherical plots of the light scattering cross-section shown in Fig. S14 represent the supplementary data for Fig. 4B. When compared with the platelet without the grating (right column), the platelet with the grating showed strong light scattering toward some specific directions.

Figure S15 shows the electric field distributions around the guanine platelet models when simulated using the FDTD software. The ripple pattern in the electric field distribution calculated for the platelet with the grating (Fig. S15A) is believed to be the origin of the directional and strong side- and back- scattering. When the angle of incidence was 60° (Fig. S15B), we observed the formation of a diffraction pattern on the transmission side that was organized different manner to the scattering on the reflection side.

The spherical plots of the light scattering cross section shown in Fig. S16 represent the supplementary data for Fig. 4C. The plots show the diversity of the anisotropic light scattering although regular reflection also exists at the same angle.

Figure S17 also shows supplementary data for Fig. 4C. The changes in the electric field distributions around the model platelet during and after passage of the incident light through the platelet are presented. We can see that an oscillation in the electric field intensity along the platelet remains. It appears that the remaining light wave emerged from the two narrow edges of the platelets. This phenomenon may possibly be related to the whispering-gallery-mode (WGM) oscillation of light (29).

The results presented in Fig. 4C were studied by changing the model from the platelet with-nano-holes to a plane mono-platelet, as illustrated in Fig. S18A. When compared with the diffraction pattern obtained with the nano-holes, the light diffraction produced by the mono-platelet was less remarkable; however, a similar diffraction pattern remained. The electric field distributions (Fig. S18B) of the mono-platelet also produced the previously observed WGM-like behavior.

These simulation results (Figs. S17, S18) are consistent with our previous findings, where light emission was detected from the edges of guanine platelet when the platelet was oriented under magnetic fields of up to 5 T (30). In a future study, we it will be necessary to distinguish between this delayed light propagation from the platelet edges and that from the light diffraction caused by the nano-holes, in the on-chip projected images.

With regard to the formation of the colored variant images on the sensor chip, additional FDTD simulations were performed to support the discussion in the main text (Fig. S19 ~ S22). The new model (Fig. S19A) contains a linear grating, but this is the only difference from the previous model. The properties of the light source (Fig. S19B) remain the same.

Figures. S20 and S21 show the results of FDTD simulations of the three stacked guanine platelets with nano holes. The spacings between the platelets were 570 nm and 1100 nm- 1150 nm for the models in Figs. S20 and S21, respectively. In the spherical cross-section plots, distinct light scattering peaks were indicated using asterisk (*). In the former case (Fig. S20), the diffraction peaks of light denoted in this way from 480 nm- 500 nm and the peaks from 570 nm-630 nm were separated on the transmitted-diffraction side. A distinct scattered light peak appeared separately from the regular reflection on the reflection side. Incident light at 450 nm caused strong light scattering, causing the light scattering beams with green to red colors to overlap with the blue beams. Expansion of the spacing (Fig. S21) modified the allocation of these distinct light scattering beams.

Figure S22 shows the case of the model with disordered platelet alignment. The platelet located at the center of the stack is not in parallel with the other platelets. It was then apparent that the strong light scattering at 450 nm disappeared, and the color specificity in the light scattering beams also decreased.

These data support the speculation in the main text regarding the mechanism for production of the colored variant images.

### Supplementary Text

Grating patterns on fish guanine platelet- technical comments

Even when the electron microscopes (i.e., SEM and TEM) were used, it was necessary to treat the guanine platelet surface to find the grating structures on the guanine platelets, or to find a platelet with a degraded surface (25). In this study, we successfully observed the gratings using an optical microscope. Careful tracking of the movie of the platelet to detect slight changes in its inclination was important for successful observation of the gratings.

The biogenic guanine platelets obtained from the fish were a mixture of platelets with gratings and plane “perfect” platelets. To date, the mechanism and the timing for formation of the gating on the platelet have not been clarified. We were stocking the guanine platelets from the fish in distilled water containing 0.01% sodium azide at 4 °C. Further inspections will be carried out to determine whether additional modifications occur on the platelet during the preservation period. However, the grating structures were also found in fresh samples from the Japanese anchovy. Other types of seawater fish such as the sea bream also had guanine platelets that showed grating textures.

Previous studies of the crystal structures and morphologies of the fish guanine platelets (28, 31-33, except 25) did not actually refer to any evidence for the existence of these grating structures. As mentioned in the main text and in the Materials and Methods section, a plane platelet without nano-holes can also form a strong light scattering beam on the image sensor array. However, the gratings on the thin platelets definitely enhance the intensities of the scattering peaks.

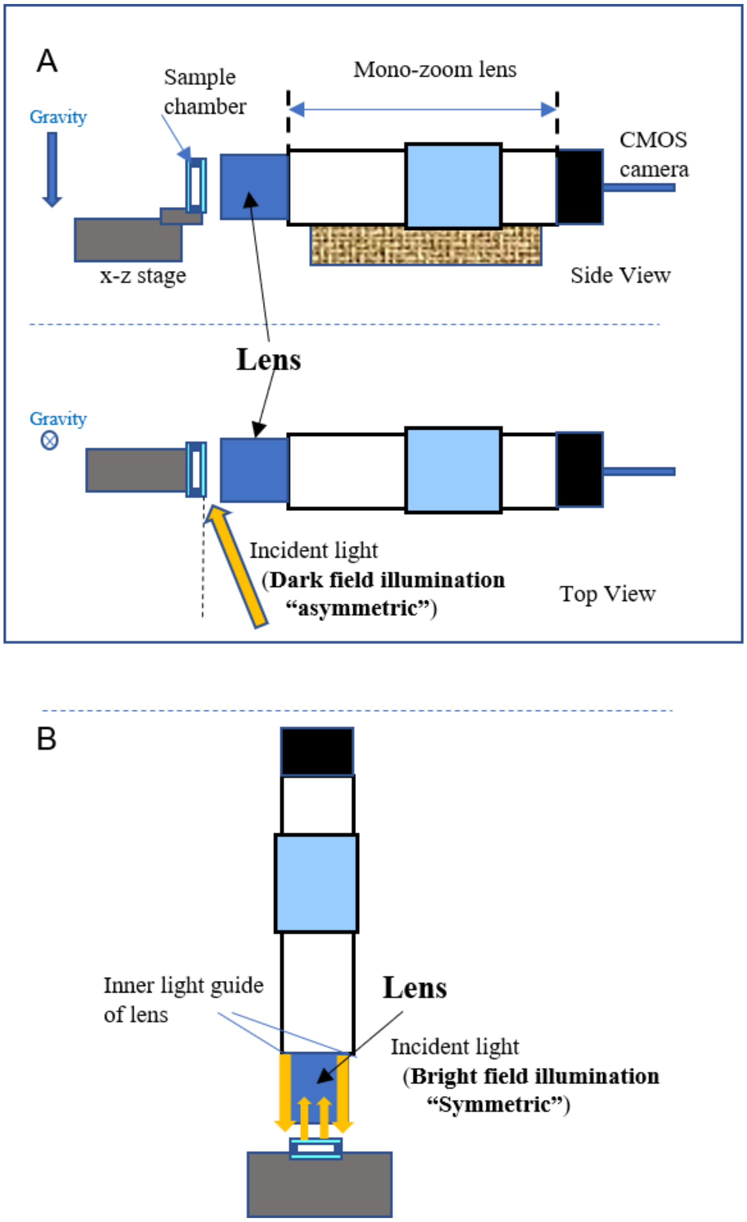


**Fig. S1.**

Microscopic observation system. (A) Asymmetric illumination from a specific direction with a separate illumination unit (optical fiber-guided light). The optical axis of the lens was set horizontally. (B) Symmetrical illumination using a light guide inside the lens.

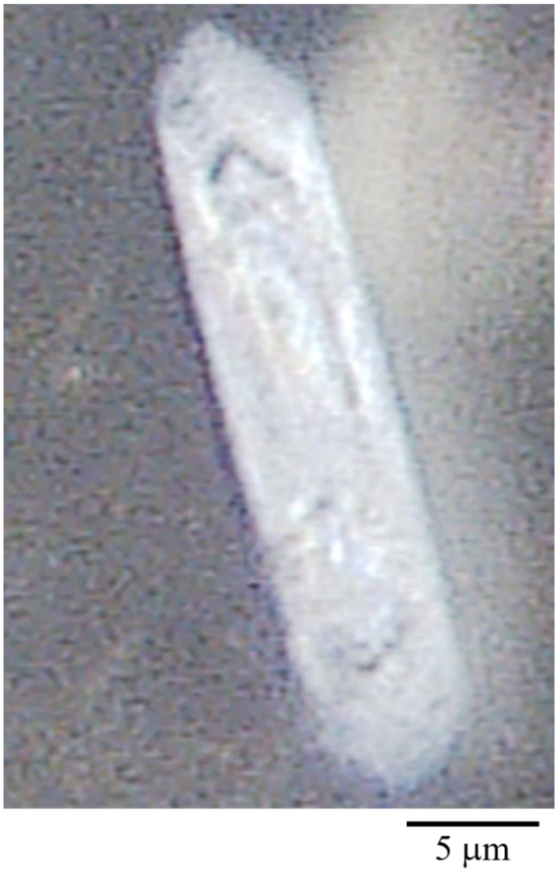


**Fig. S2.**

Image of a guanine platelet (separated from a goldfish scale) that was observed under symmetrical illumination using the light guide inside the lens.

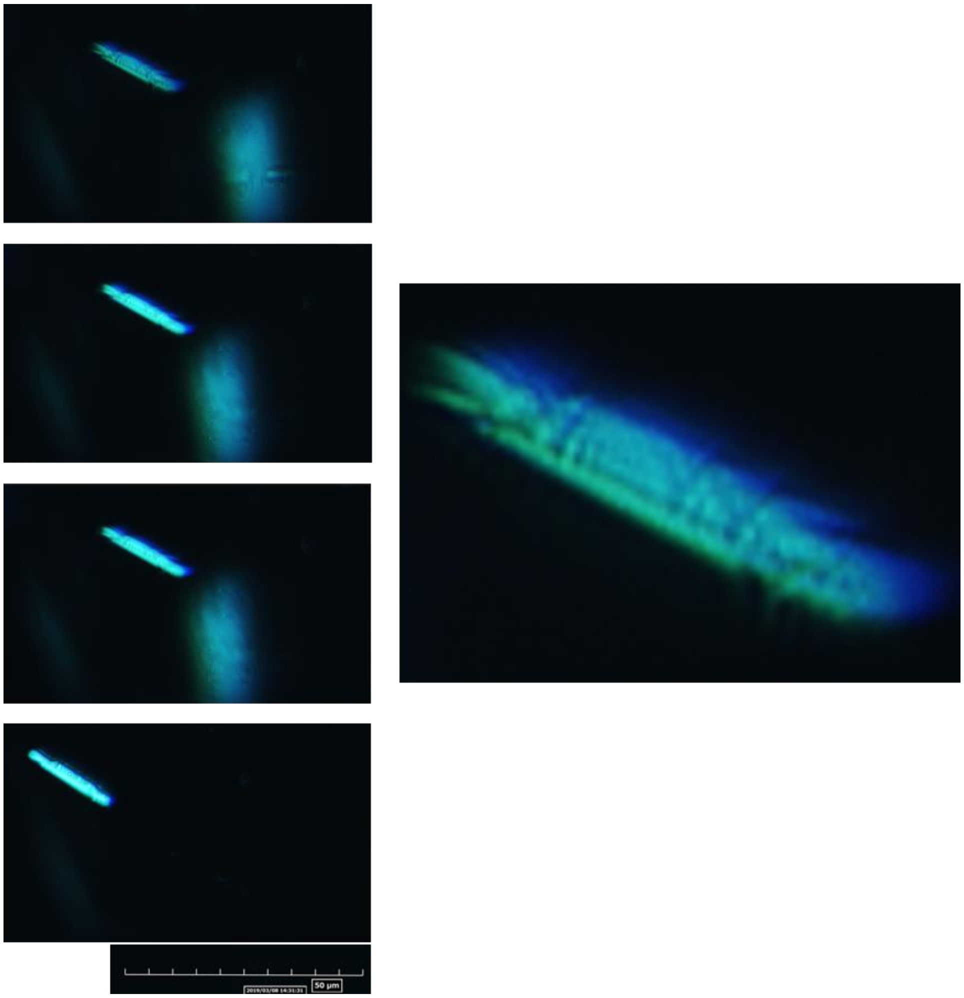


**Fig. S3.**

Images of a guanine platelet (separated from a goldfish scale) observed under asymmetric illumination using a separate illumination unit (optical fiber-guided light).

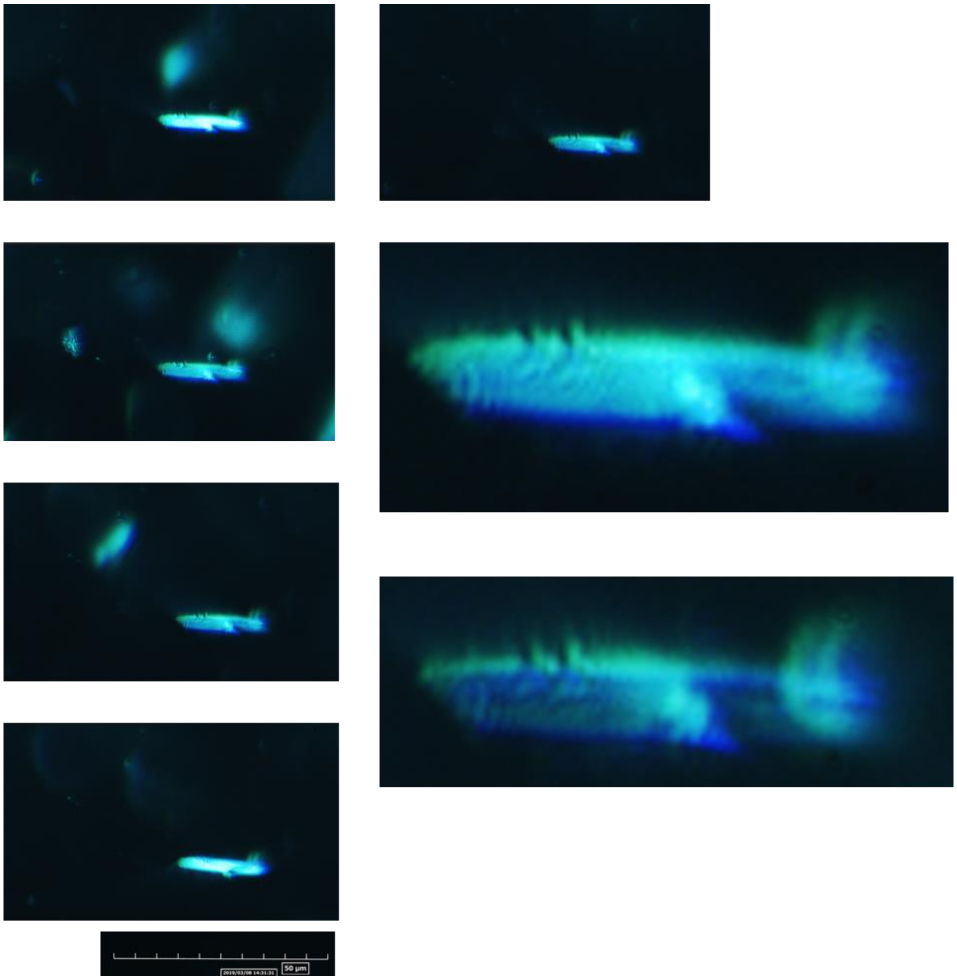


**Fig. S4.**

Further example images of the guanine platelet (separated from a goldfish scale) observed under asymmetric illumination using a separate illumination unit (optical fiber-guided light).

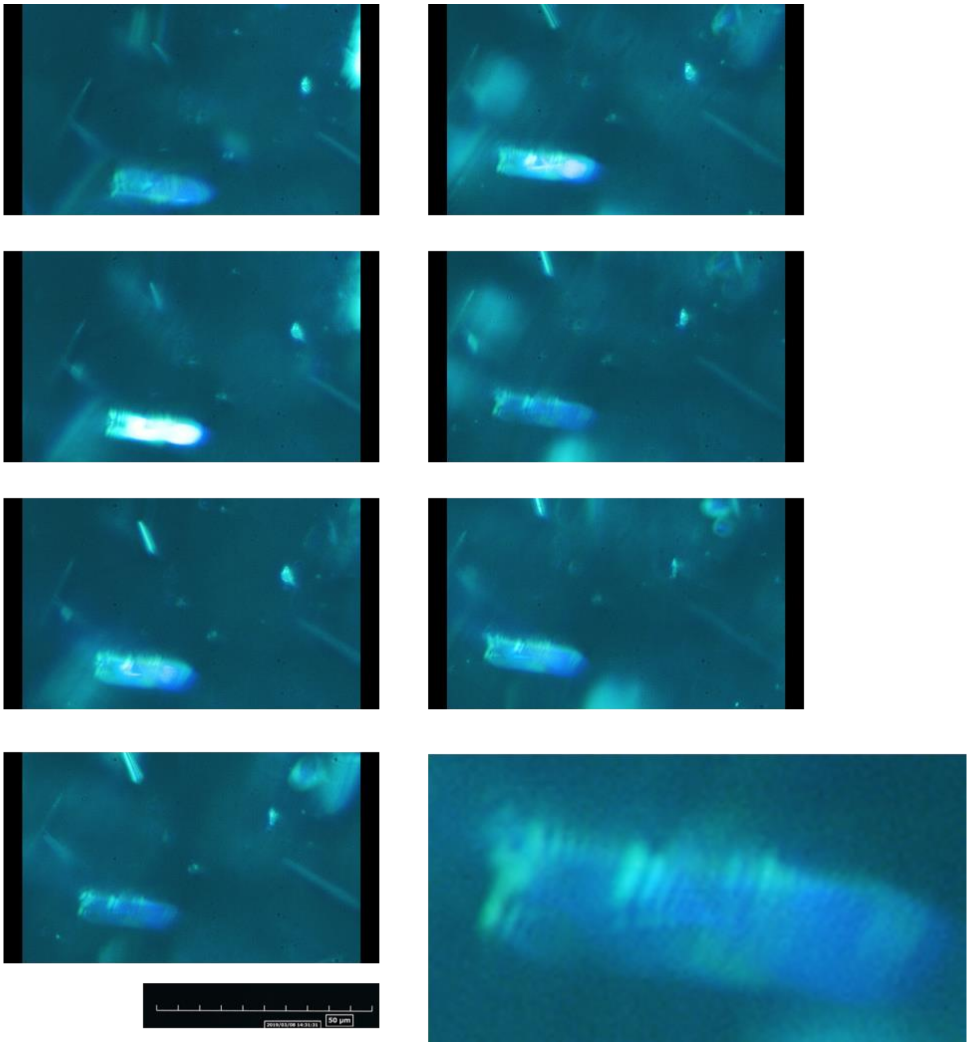


**Fig. S5.**

Images of the guanine platelet (separated from a Japanese anchovy) observed under asymmetric illumination using a separate illumination unit (optical fiber-guided light).

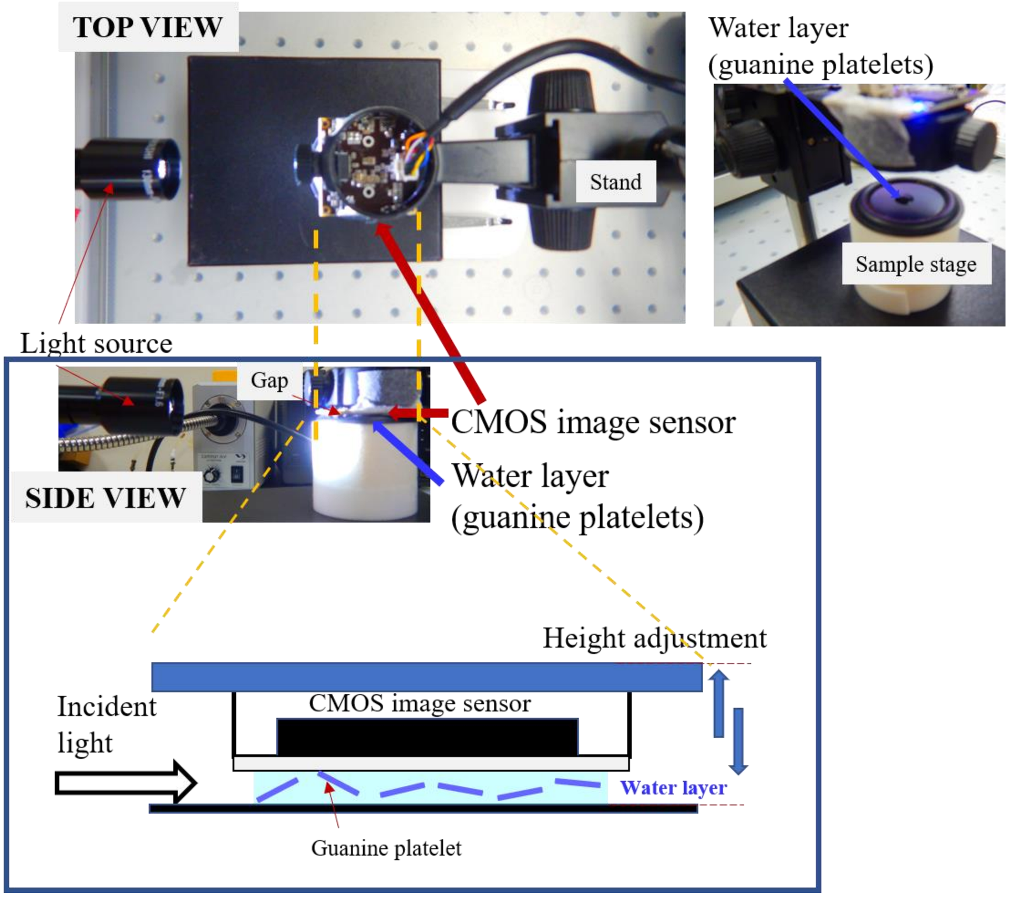


**Fig. S6.**
Top and side views of the on-chip imaging system used for imaging of the fish guanine platelets in the thin water layer between the CMOS image sensor and the light shielding plate. The water layer thickness can be adjusted by varying the height of the image sensor.

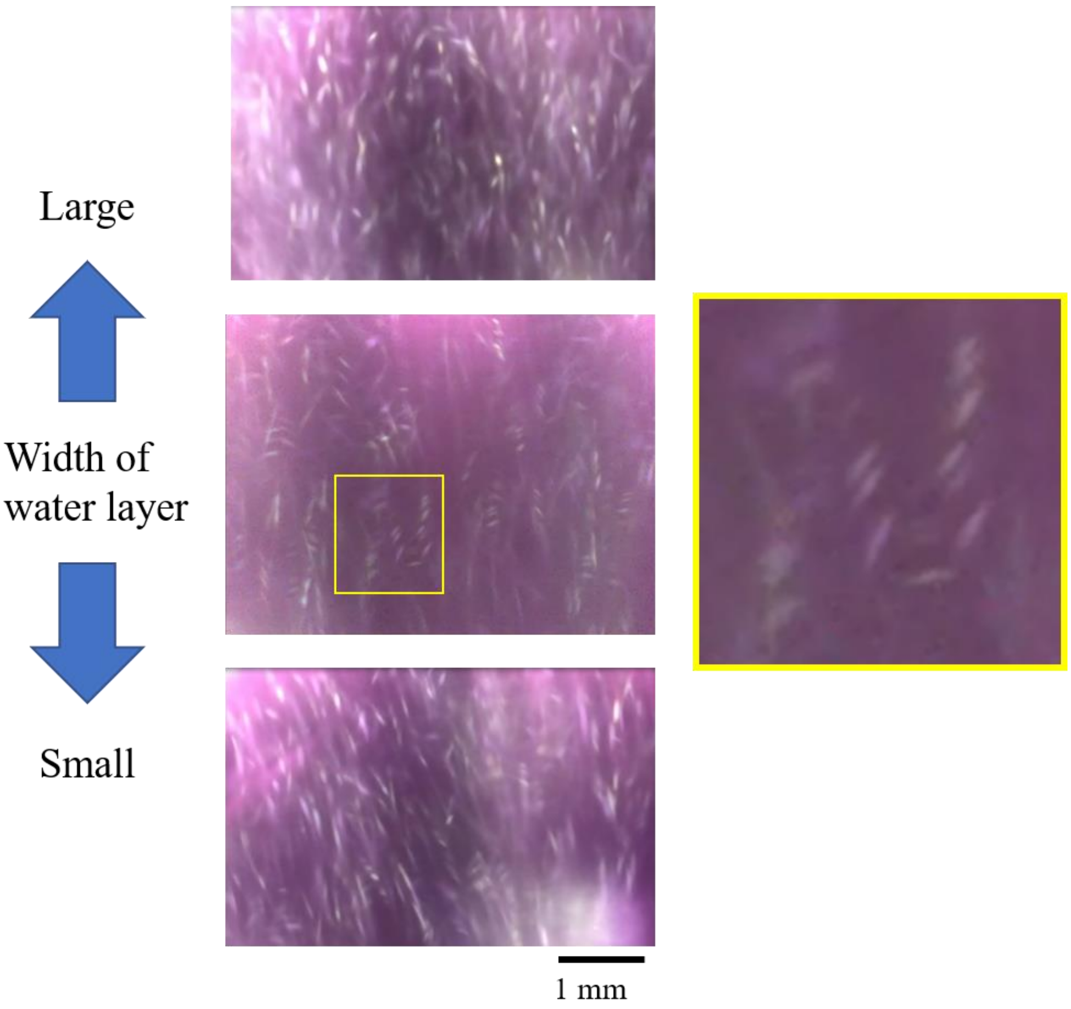


**Fig. S7.**

Changes in the pattern of images projected on the CMOS image sensor that occurred when the water layer width was either extended or shortened.

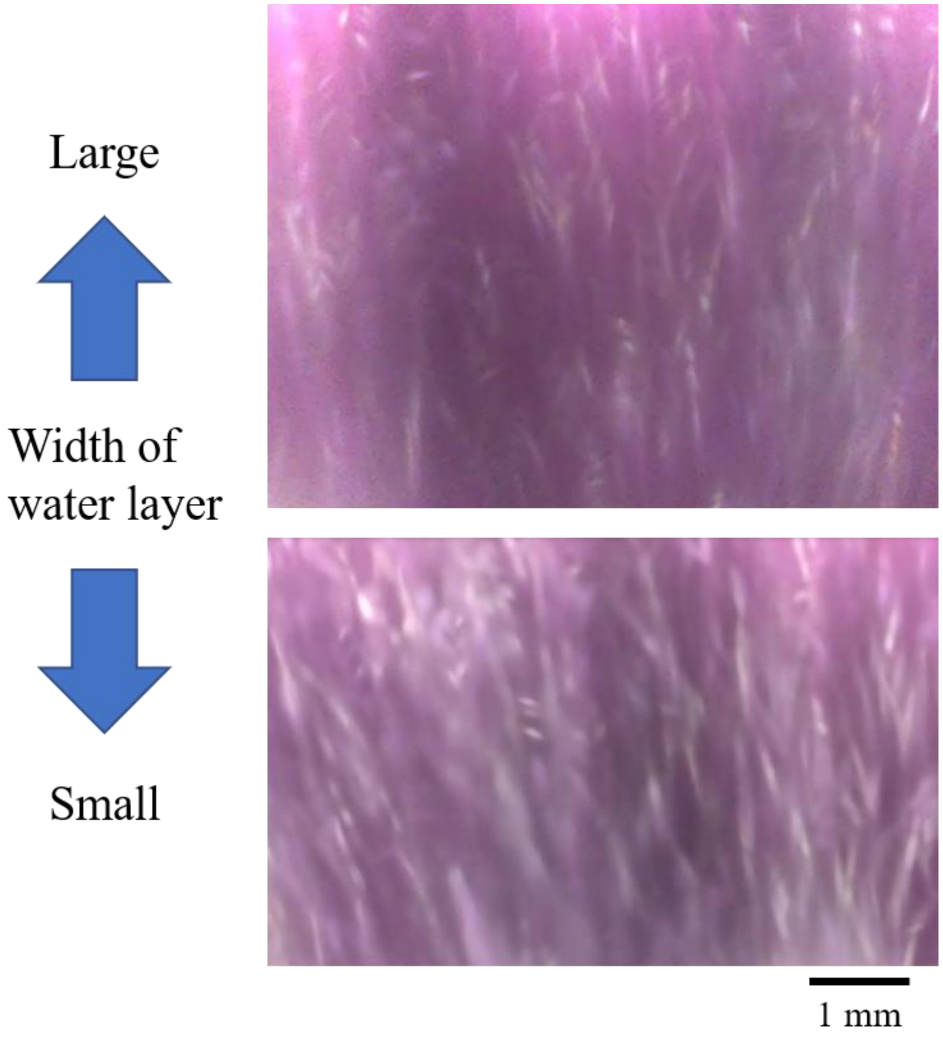


**Fig. S8.**

Another example of the changes in the pattern of images projected on the CMOS image sensor that occurred when the water layer width was either extended or shortened.

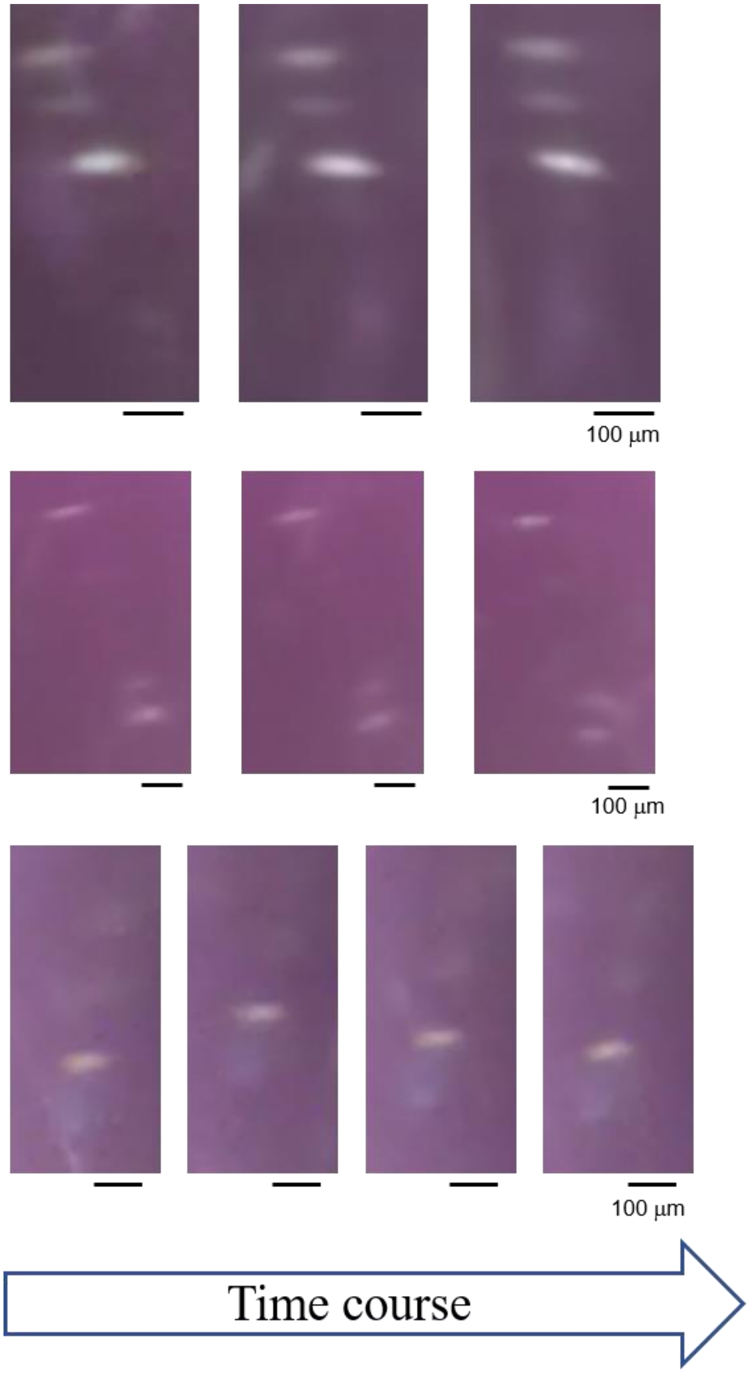


**Fig. S9.**

Time courses of the movement of the bar-code-type projected images that show evidence of the bar-code images moving together.

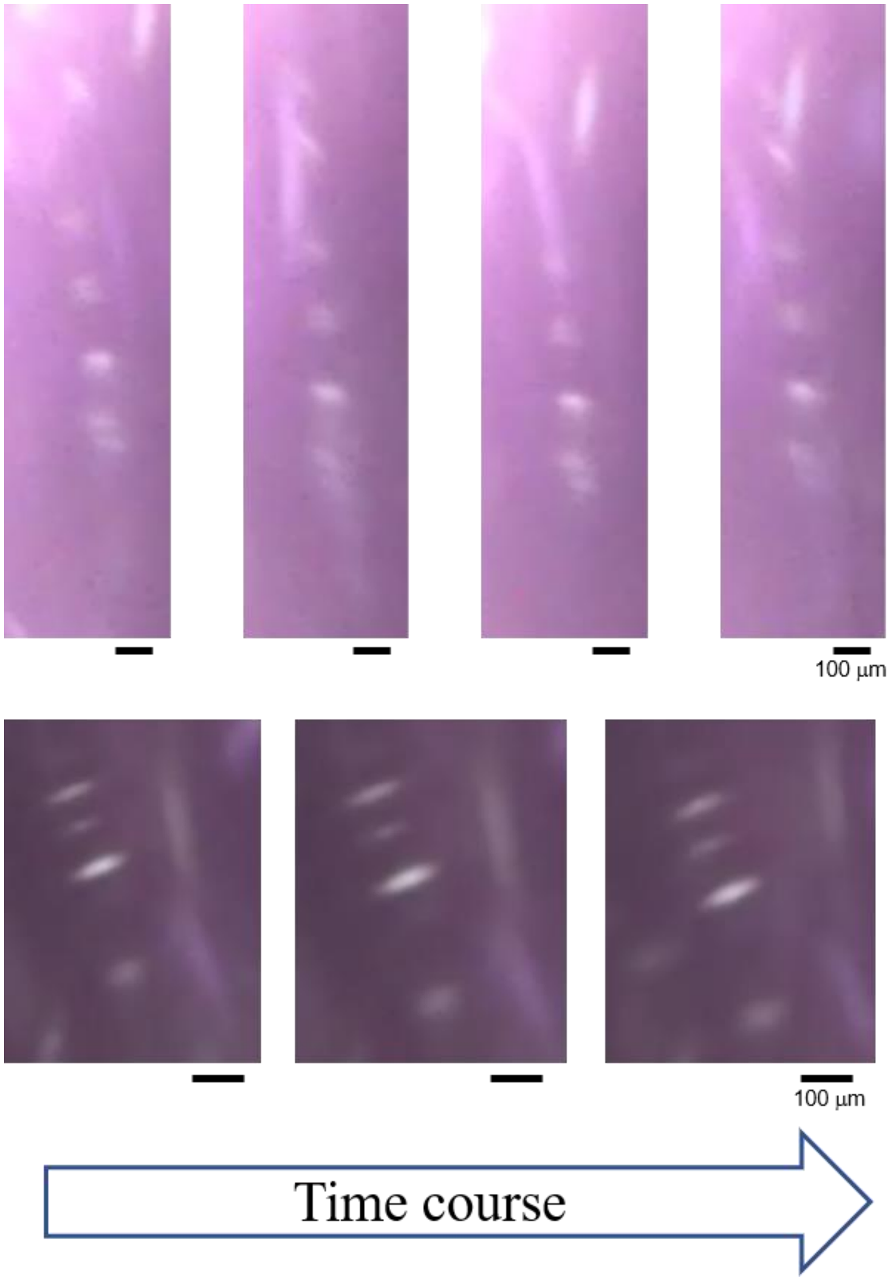


**Fig. S10.**

Other examples of the time courses of the movement of the bar-code-type projected images with seven bars (top) and four bars (bottom).

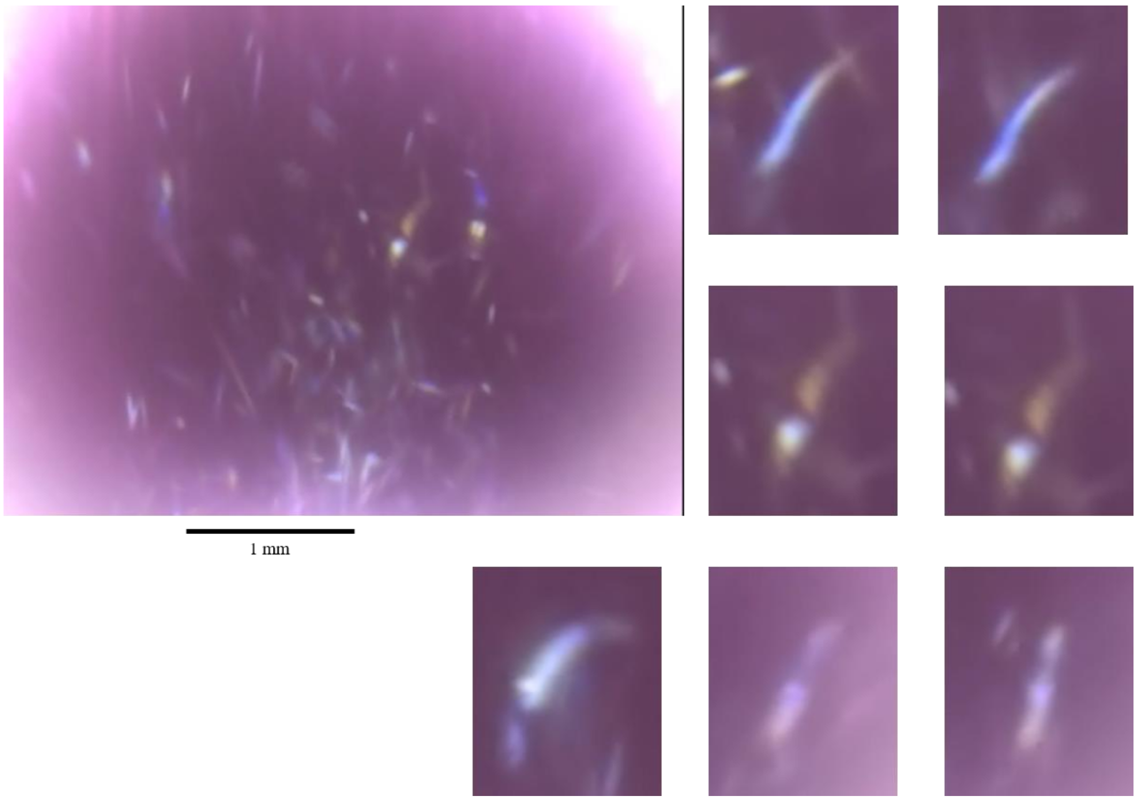


**Fig. S11.**
Colored variants of the column-type images that were projected directly onto the CMOS sensor.

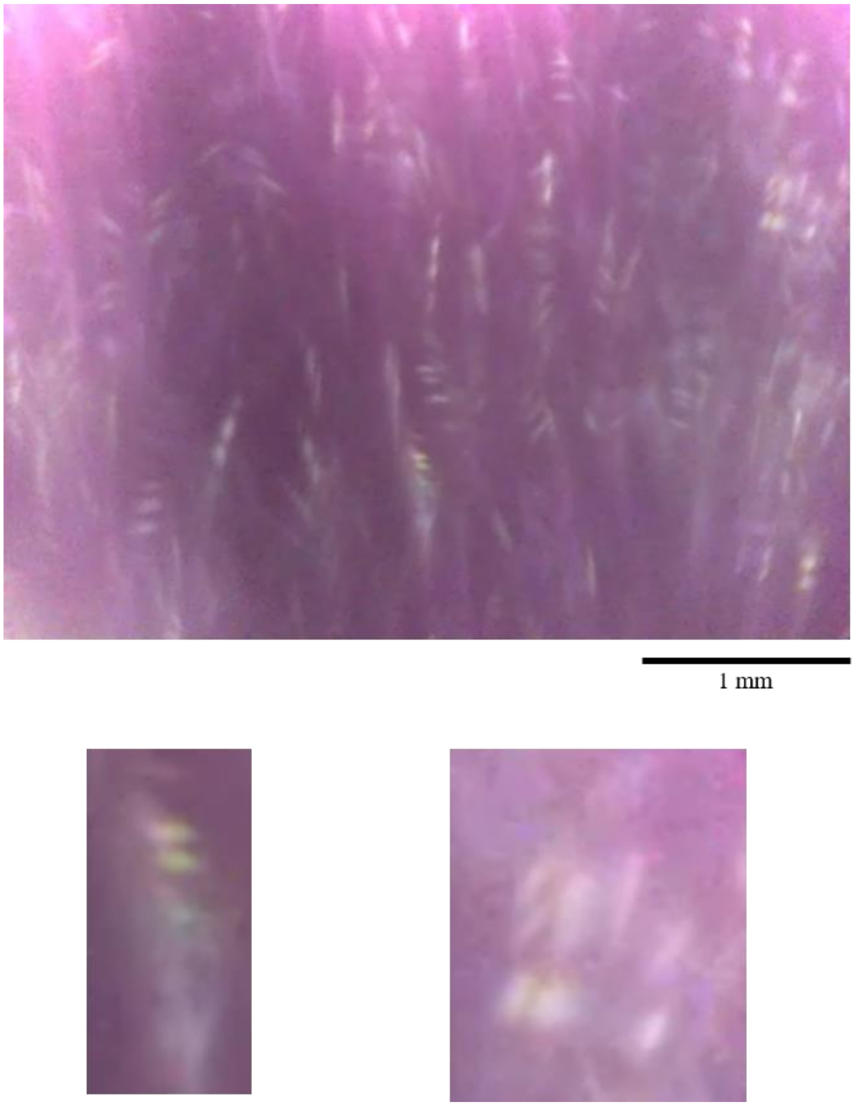


**Fig. S12.**

Bar-code images projected on the entire area of the CMOS sensor array. Variants, such as a colored pattern of mono-platelets (bottom -left) and a pattern composed of connected platelets are included.

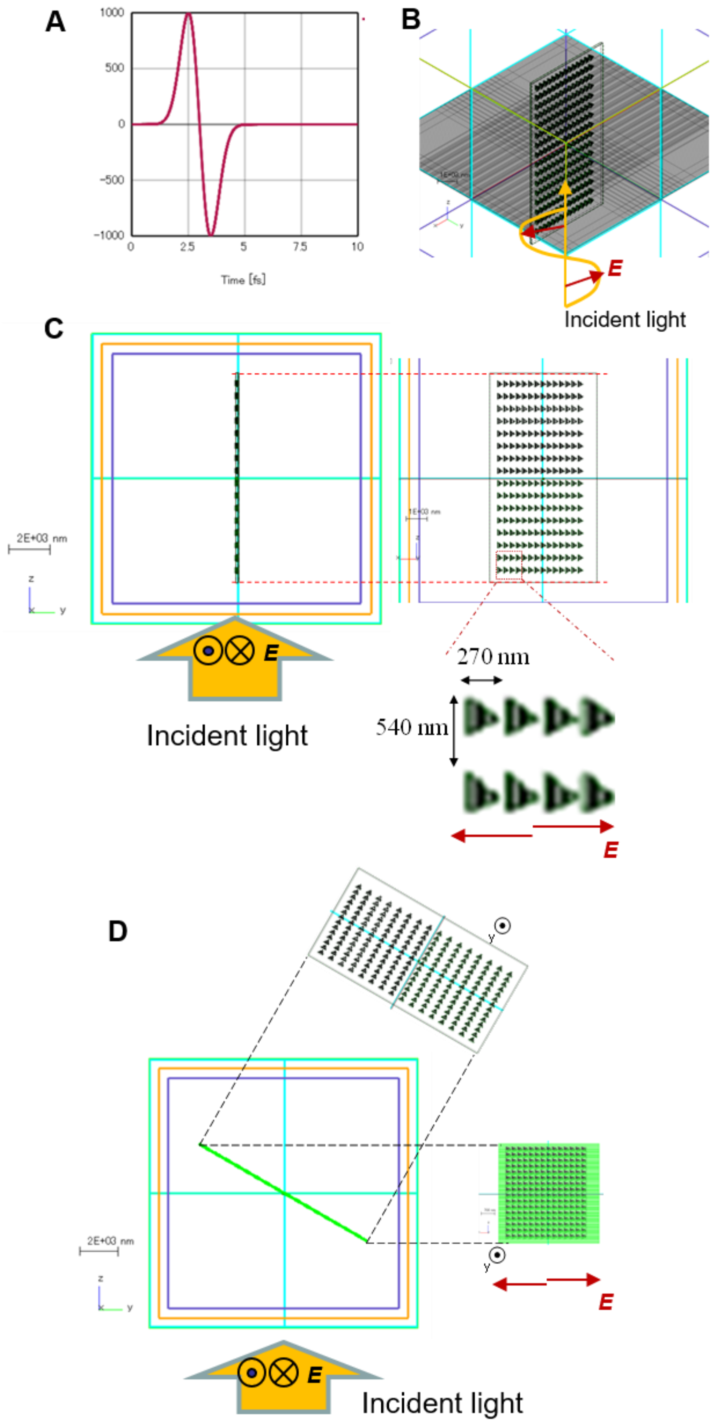


**Fig. S13.**

Model of guanine platelet with nano-hole gratings for the FDTD simulations. (A) Pattern of the incident pulsed light. (B) Relationship between platelet orientation and the electric field direction of the (polarized) incident light. (C) Side and front views of the model of the platelet oriented in parallel with the incident light. (D) Side and front views of the model of the platelet when tilted by 60° with respect to the incident light.

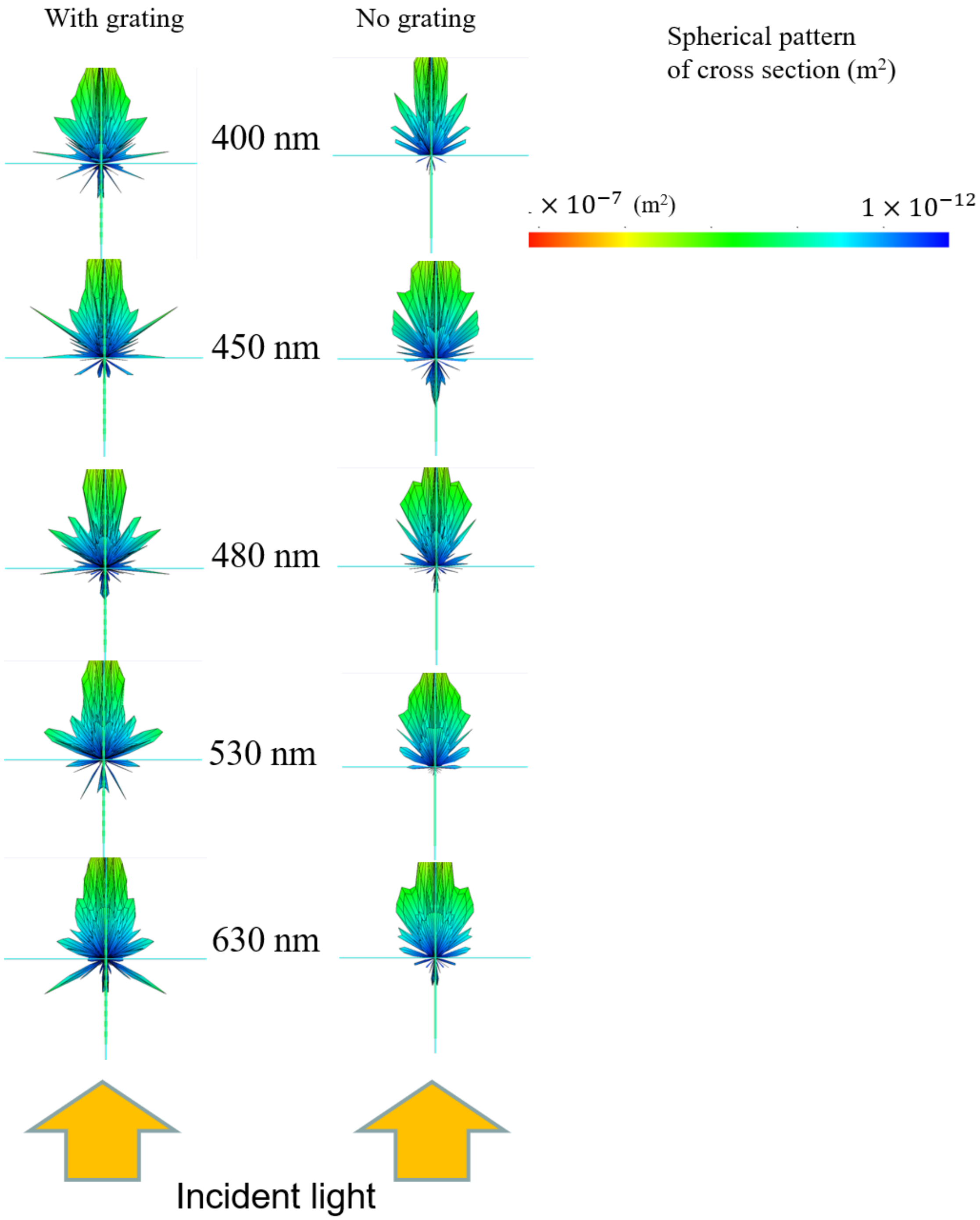


**Fig. S14.**
Supplementary data for Fig. 4B.

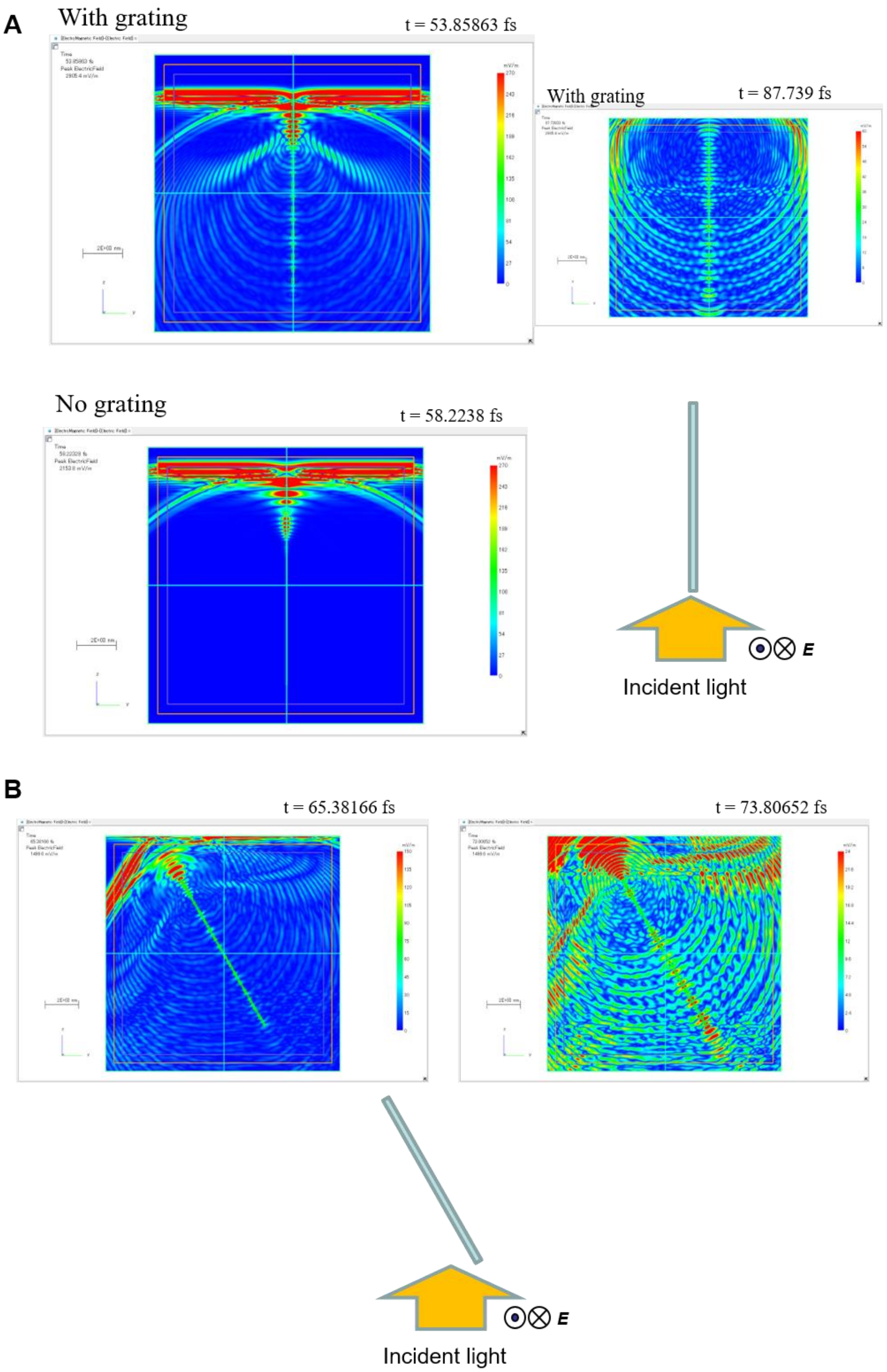


**Fig. S15.**

Electric field distributions around the guanine platelet models when simulated using the FDTD method. The incident light (large orange-colored arrow) and platelet tilting conditions are also illustrated. (A) Comparison of light scattering patterns on platelets with and without the nano holes. The platelets are oriented in parallel with the incident light. (B) Time course of changes in the electric field distributions when the angle of incidence with respect to the platelet was 60°.

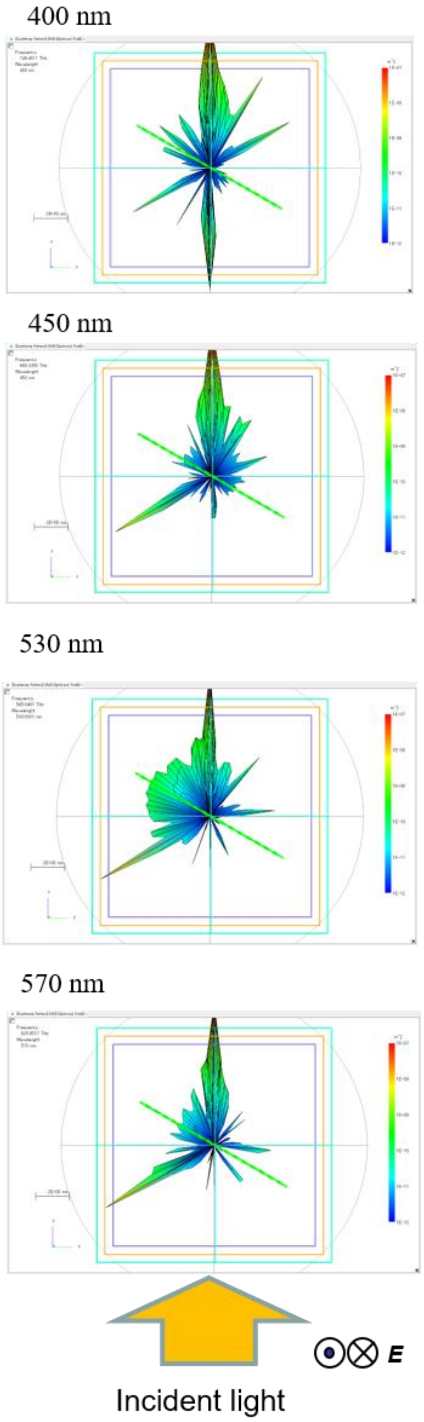


**Fig. S16.**
Supplementary data for Fig. 4C.

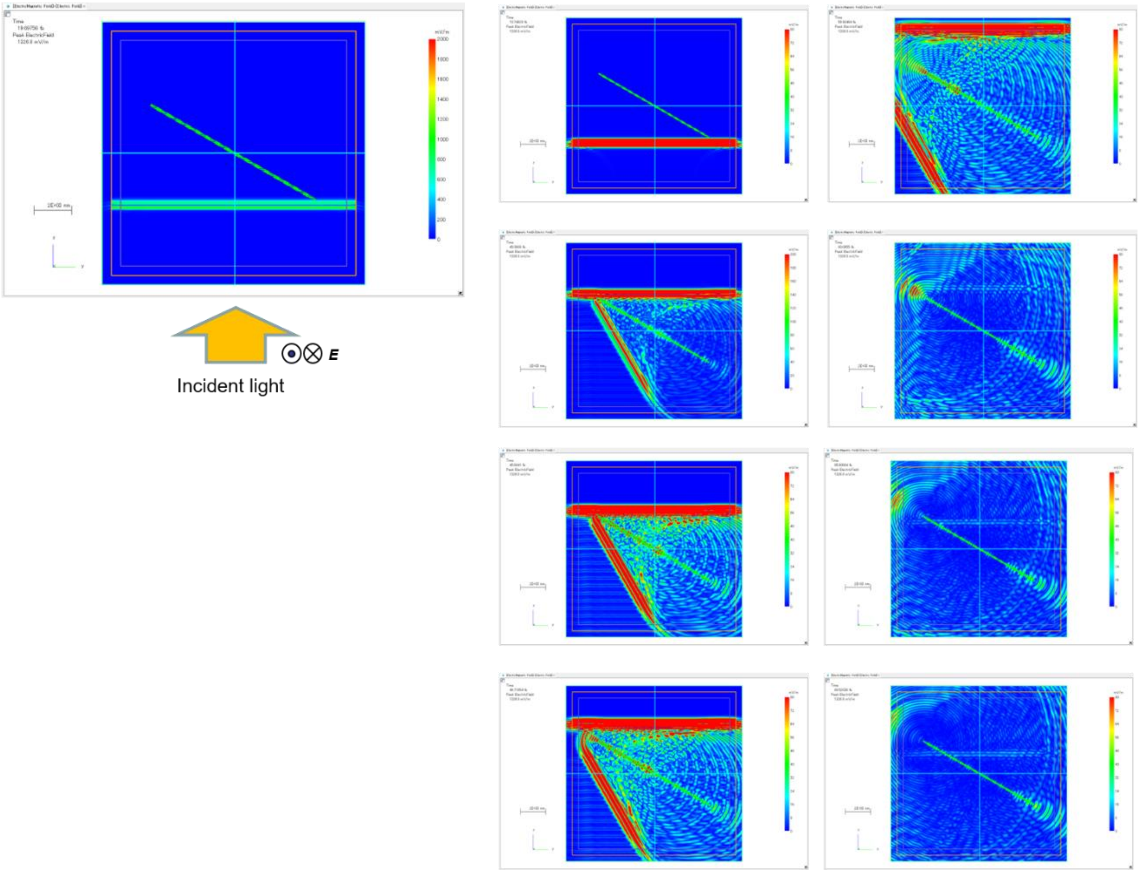


**Fig. S17.**

Supplementary data for Fig. 4C. Time courses of the changes in the electric field distributions around the model platelet when light with an angle of incidence of 60° with respect to the platelet passed through the platelet.

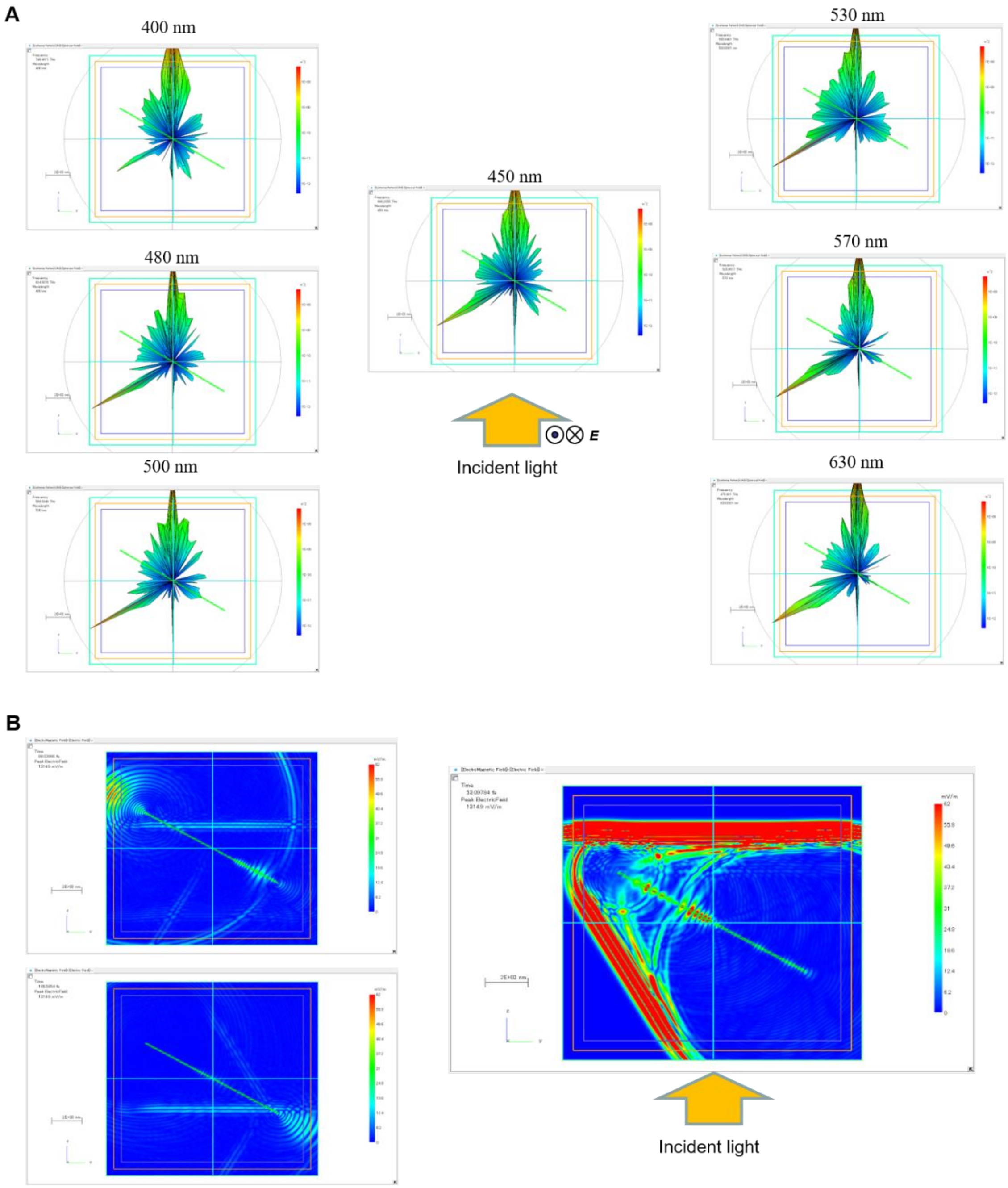


**Fig. S18.**

Supplementary data for Fig. 4C. FDTD simulation result for the guanine platelet without nano holes are shown. The angle of incidence with respect to the platelet was 60°. (A) Spherical plots of the scattered light cross-sections over the range from 400 nm to 630 nm. (B) Electric field distributions around the model platelet after the incident light had passed through the platelet.

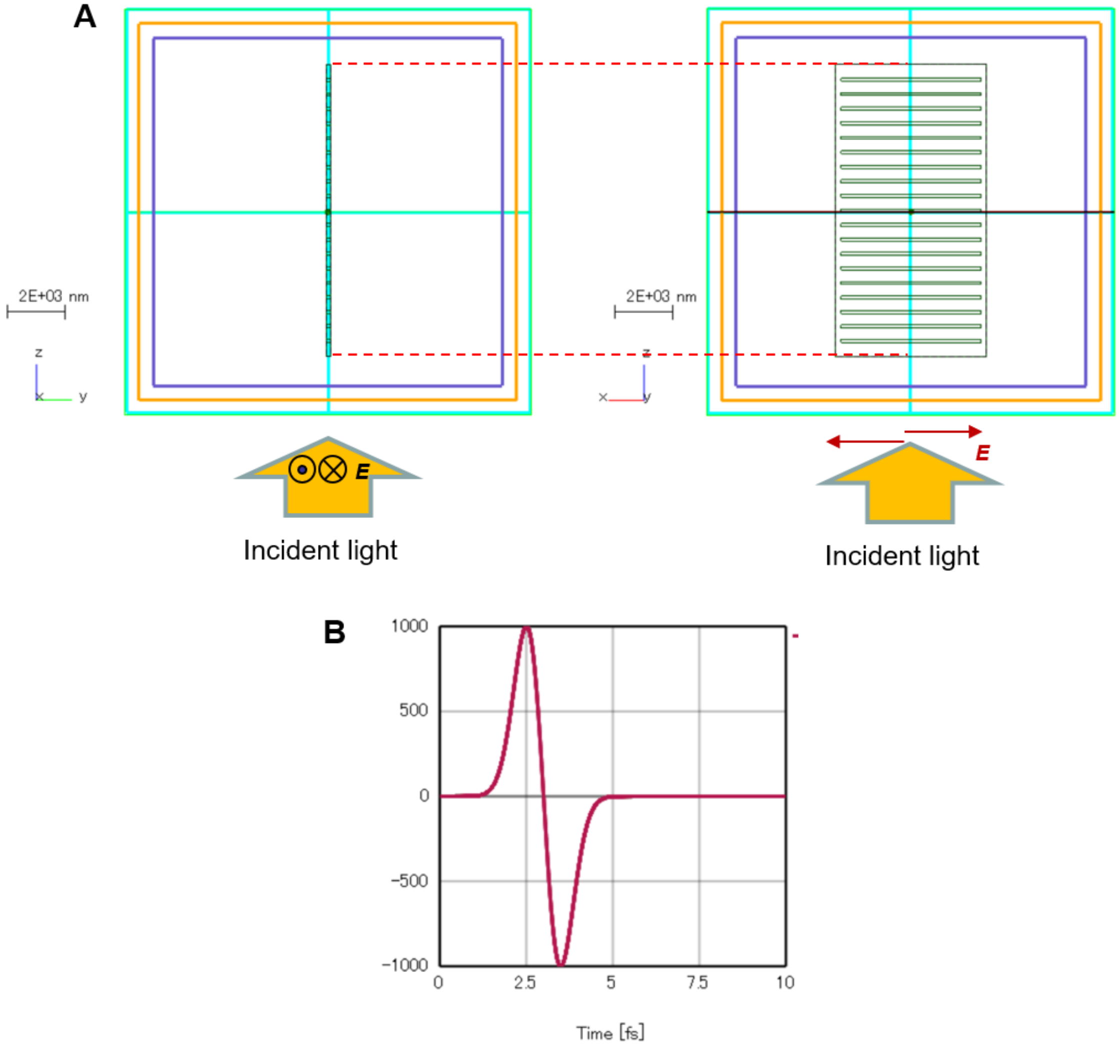


**Fig. S19.**
Model of the guanine platelets with linear gratings for the FDTD simulations. (A) Side and front views of the model of the platelet oriented in parallel with the incident light. (B) Pattern of the incident pulsed light.

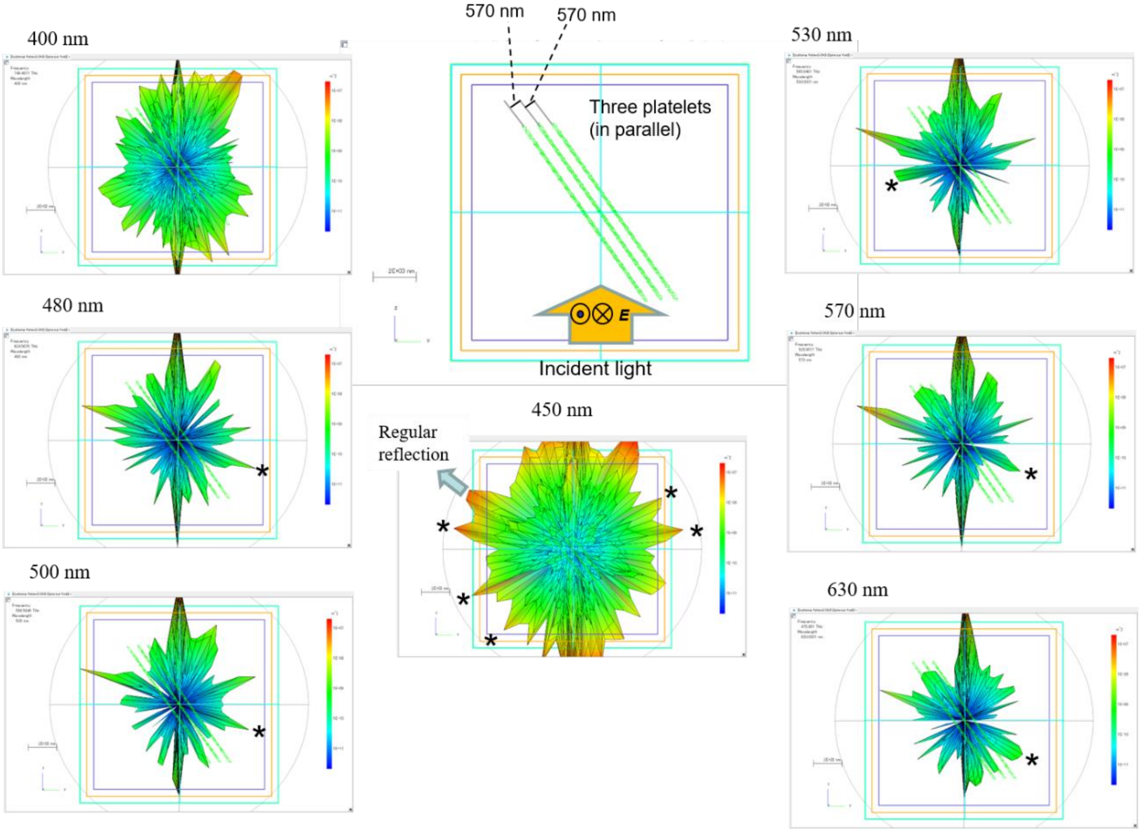


**Fig. S20.**

FDTD simulation results for three stacked guanine platelets with nano holes. The angle of incidence with respect to the platelet was 60°. Spherical plots of the scattered light cross-section over the range from 400 nm to 630 nm are shown.

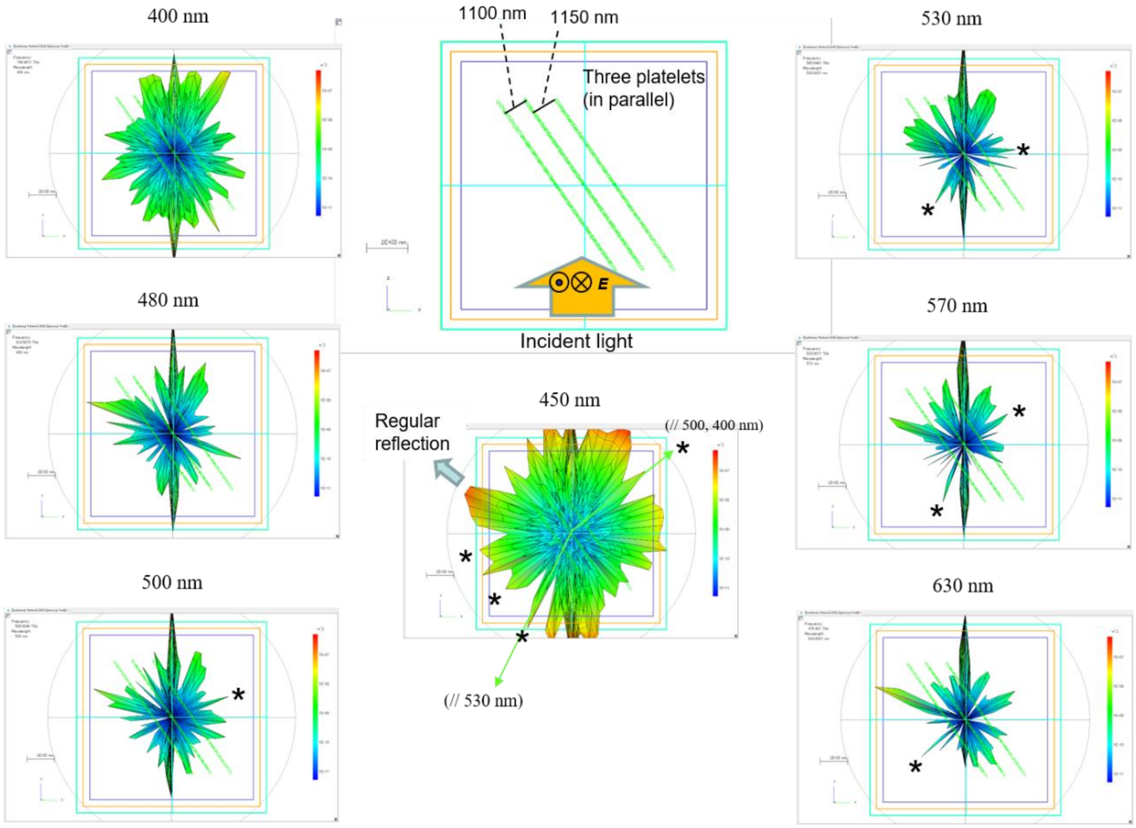


**Fig. S21.**

FDTD simulation results for three stacked guanine platelets with nano holes. The spacing between the platelets is greater than that of the model in Fig. S20. The angle of incidence with respect to the platelet was 60°. Spherical plots of the scattered light cross-sections over the range from 400 nm to 630 nm are shown.

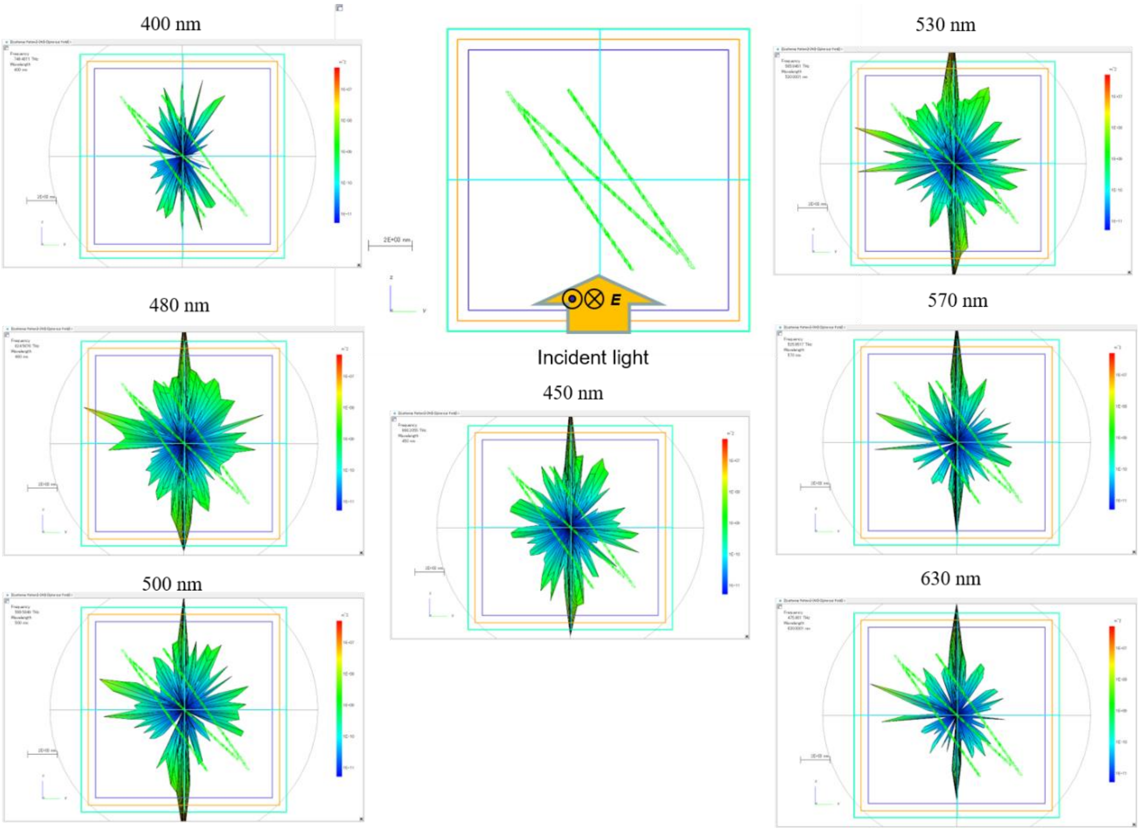


**Fig. S22.**

FDTD simulation results for the three stacked guanine platelets with nano holes. The platelet located at the center of the stack is not in parallel with the other platelets, as shown in the center-top panel. Spherical plots of the scattered light cross-sections over the range from 400 nm to 630 nm are shown.

**Movie S1.**

Movie of on-chip light scattering imaging of the tiny platelets from goldfish; an image of these platelets is shown in Fig. 2E in the main text. Bar-code-type projected images are presented.

**Movie S2.**

Movie of the colored patterns of light scattered by the guanine platelets from the Japanese anchovy; images of these platelets are shown in Fig. 3 in the main text.
